# Genomic history and ecology of the geographic spread of rice

**DOI:** 10.1101/748178

**Authors:** Rafal M. Gutaker, Simon C. Groen, Emily S. Bellis, Jae Y. Choi, Inês S. Pires, R. Kyle Bocinsky, Emma R. Slayton, Olivia Wilkins, Cristina C. Castillo, Sónia Negrão, M. Margarida Oliveira, Dorian Q. Fuller, Jade A. d’Alpoim Guedes, Jesse R. Lasky, Michael D. Purugganan

## Abstract

Rice (*Oryza sativa*) is one of the world’s most important food crops. We reconstruct the history of rice dispersal in Asia using whole-genome sequences of >1,400 landraces, coupled with geographic, environmental, archaeobotanical and paleoclimate data. We also identify extrinsic factors that impact genome diversity, with temperature a leading abiotic factor. Originating ∼9,000 years ago in the Yangtze Valley, rice diversified into temperate and tropical japonica during a global cooling event ∼4,200 years ago. Soon after, tropical rice reached Southeast Asia, where it rapidly diversified starting ∼2,500 yBP. The history of indica rice dispersal appears more complicated, moving into China ∼2,000 yBP. Reconstructing the dispersal history of rice and its climatic correlates may help identify genetic adaptation associated with the spread of a key domesticated species.

**One sentence summary:** We reconstructed the ancient dispersal of rice in Asia and identified extrinsic factors that impact its genomic diversity.

---

Rice (*Oryza sativa* L.) is a major staple crop, providing > 20% of calories for more than half of the human population. Domesticated rice encompasses genetically distinct populations grown in sympatry, including major subgroups japonica and indica (sometimes recognized as subspecies), as well as geographically more restricted *circum*-aus, and *circum*-basmati rices (*1, 2*). It is mainly cultivated in monsoon Asia, but rice is distributed across a wide latitudinal range, spanning tropical and temperate zones of Asia, likely requiring local water, temperature and photoperiod adaptation. Rice is grown in lowland ecosystems under paddy, deepwater, or seasonal flood conditions, as well as in upland rainfed areas (*3*).

Archaeological evidence (*4–6*) indicates that cultivation of japonica rice began ∼9,000 years before present (yBP) in the lower Yangtze Valley, while proto-indica rice cultivation started >5,000 yBP in the lower Ganges valley (*7*). Archaeological (*8*) and most population genetic analyses (*9–11*) suggest that important domestication alleles have a single origin in japonica rice in East Asia. The spread of japonica to South Asia ∼4,000 years ago led to introgression of domestication alleles into proto-indica or local *O. nivara* populations and the emergence of indica rice (*9–11*). From the Yangtze and Ganges Valleys, respectively, japonica and indica dispersed across much of Asia over the last 5 millennia, providing sustenance for emerging Neolithic communities in East, Southeast and South Asia (*12*).

Archaeological data shows the general directionality of rice dispersal (*7, 13*); the details of dispersal routes, times, and the environmental forces that shaped dispersal patterns, however, remain unknown. Here, we undertake population genomic analyses to examine environmental factors associated with the geographic distribution of rice diversity, and reconstruct the ancient dispersal of rice in Asia. Together with archaeobotanical, paleoclimatic and historical data, genomic data allows a robust reconstruction of the dispersal history of *Oryza sativa*.

## Structure of rice genomic diversity

We obtained whole genome re-sequencing data from rice landraces/traditional varieties across a wide geographical distribution in Asia. Our sample set includes 1,265 samples from the Rice 3K Genome Project (*1*) and additional 178 landraces sequenced for this study (Supplementary Table 1); the panel consists of 833 indica, 372 japonica, 165 *circum*-aus, 42 *circum*-basmati, and 31 unclassified samples. We identified ∼9.78 million single nucleotide polymorphisms (SNPs) with 9.63x mean coverage (s.d. = 5.03), which we used in subsequent analyses (Supplementary Fig. 1).

Analysis of molecular variance (AMOVA) indicated that subspecies affiliation explained >36% of the total variation [AMOVA, permutation *P* < 0.001](*14*), congruent with results from multidimensional scaling (MDS) of genomic distances (Supplementary Fig. 2a). Only japonica and indica have wide geographic distributions (Fig. 1 a and b; Supplementary Fig. 3), and AMOVA of these two subspecies (n=1,205) revealed that genomic variance is explained by subspecies (*r*^*2*^ = 0.32, permutation *P* < 0.001), country of origin (*r*^*2*^ = 0.11, *P* < 0.001) and their interaction (*r*^*2*^ = 0.06, *P* < 0.001). Landraces with mixed ancestry (n=154) were excluded using silhouette scores (Supplementary Fig. 2b); henceforth, we analysed these two subspecies independently.

**Figure 1:**
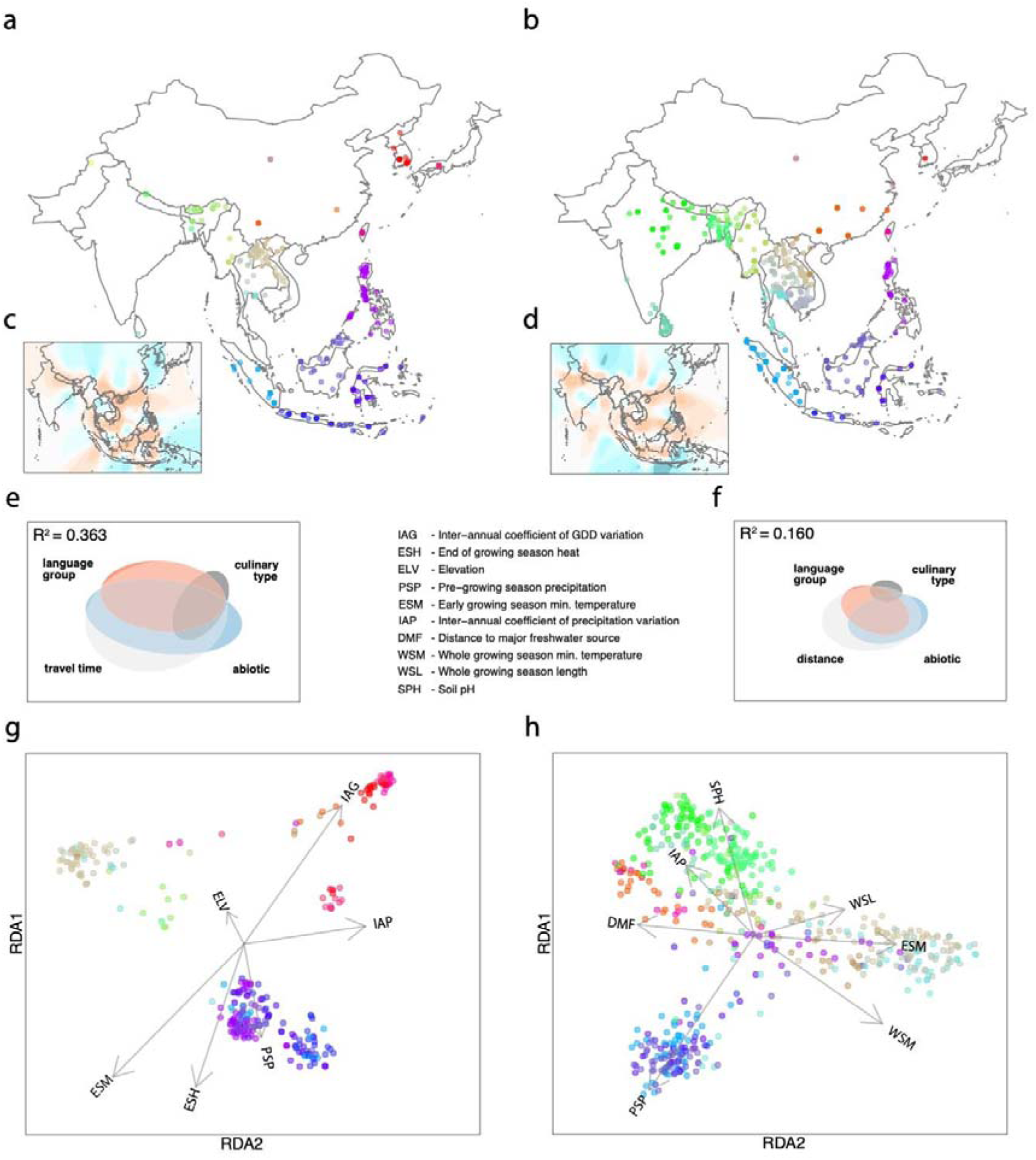
Factors underlying geographic distribution of genomic diversity in japonica and indica. Maps of collection sites for (a) japonica and (b) indica landraces used in this study. Colors represent regions of origin. (c) Japonica and (d) indica effective migration surfaces representing migration barriers (orange) and channels (cyan). (e) Japonica and (f) indica genomic diversity is best explained by a combination of four factors represented in Euler plots: travel time (migration resistance) or geographic distance, abiotic variables (temperature, moisture and soil characteristics), linguistic group, and culinary properties (stickiness). Fields of squares represent total genomic variation, while elliptic shapes represent genomic variation explained by particular factor. (g) Japonica and (h) indica genotypes projected on the first two canonical axes of redundancy analysis. Arrows represent environmental predictors (acronyms explained in the legend) that strongly correlate with a maximal proportion of linear combinations of SNPs.

We find support for isolation-by-distance (IBD) in japonica (*r*^*2*^ = 0.294, *P* < 0.001) and indica (*r*^*2*^ = 0.265, *P* < 0.001) [Supplementary Fig. 4]. Geographic distance explains genetic distance much less in the Malay Archipelago (*i.e.* islands SE Asia) compared to mainland Asia, suggesting a stronger effect of local migration barriers on island IBD (Supplementary Fig. 5). Effective migration surfaces (*15*) identified geographic barriers for dispersal over the Hengduan Mountains which separate China from South/Southeast Asia, and the South China Sea which reduces movement between Borneo/Philippines and mainland Southeast Asia (Fig. 1c and d; Supplementary Fig. 6). For human-dispersed species such as crops, genetic distances may correlate better with travel resistance, meant to capture cost-effective migration by humans. An isolation-by-resistance (IBR) model, using estimated human-associated land and marine travel times (*16*), is a better explanation than the IBD model for japonica landrace genetic distances based on Akaike Information Criterion (island ΔAIC = -34, mainland ΔAIC = -17), but not for indica (island ΔAIC = +51, mainland ΔAIC = +611)[Supplementary Fig. 5)].

## Factors associated with spatial genomic structure

We used redundancy analysis (RDA) to partition genomic variance (*17*) associated with 22 different variables that include climatic and edaphic conditions, as well as interactions with humans and wild relatives (Supplementary Table 1). We assume that while environments in localities fluctuate over time, current genome diversity may be determined both by current environment as well as long-term evolutionary history. SNP variation is better explained by our predictors for japonica (adjusted *r*^*2*^ = 0.363; Fig. 1e) than indica (adjusted *r*^*2*^ = 0.164; Fig. 1f). Associations between predictor sets and SNPs are substantially collinear with each other. For japonica and indica, travel time and geographic distance, respectively, explain most SNP variation (adjusted *r*^*2*^ = 0.326 and *r*^*2*^ = 0.146), followed by abiotic conditions, language groups, culinary properties (*i.e*., cooked grain stickiness), and genetic composition of proximal wild rice populations (Figs. 1e and f; Supplementary Fig. 7). Among abiotic variables for japonica, temperature explains the greatest portion of SNP variation (adjusted *r*^*2*^ = 0.180), followed by moisture (*r*^*2*^ = 0.086) and soil characteristics (*r*^*2*^ = 0.081). Similarly, temperature explains the most SNP variation in indica (*r*^*2*^ = 0.064), followed by soil characteristics (*r*^*2*^ = 0.038) and moisture (*r*^*2*^ = 0.036) (Supplementary Fig. 7), although these factors have weaker explanatory power in indica compared to japonica.

The first two RDA axes of environment-associated SNP variation (*18*) separated japonica landraces consistent with geography (Fig. 1g), recapitulating results using total SNP variation (Supplementary Fig. 8a). Temperate japonica landraces from northern latitudes are most strongly identified by alleles associated with high coefficient of inter-annual variation in growing degree days, and low minimum temperatures early in the growing season (Fig. 1g; Supplementary Fig. 9a). Temperate landraces from upland rainfed ecosystems are further characterized by alleles associated with inter-annual variation in precipitation.

For indica, the first two axes also grouped individuals by their geographic origins (Fig. 1h; Supplementary Fig. 8b). Similar to japonica, indica Malay Archipelago genotypes are characterized by alleles associated with high precipitation prior to the growing season. Mainland Southeast Asian genotypes are characterized by alleles associated with warm minimum growing season temperatures and presence of nearby freshwater sources (Fig. 1h; Supplementary Fig. 9b). The latter contrasts with indica from China and most of India, where irrigation is common and there is less reliance on natural water sources (*19*)(Supplementary Table 1). Finally, genotypes in South India are identified by alleles associated with inter-annual variation in precipitation.

## Discrete subpopulations within japonica and indica

We clustered landraces based on genomic distances by partitioning-around-medoids [PAM](*20*), identifying the number of discrete clusters (K) using silhouette scores (*21*) [see Methods]. This discretization procedure removed genetic gradients between subpopulations (Fig. 2a and 2d; Supplementary Figs. 10 and 11). We compared PAM clusters to those from the ADMIXTURE algorithm (*22*). Silhouette filtering removed individuals with spurious subpopulation assignments (Supplementary Figs. 12 and 13). In general, the clustering fit using silhouette scores is greater for japonica than indica (Supplementary Fig. 14). We find consistently higher F_ST_ values among japonica subpopulations (Supplementary Fig. 15), suggesting less migration compared to indica. Finally, subpopulations of both subspecies clearly correspond with geography (Fig. 2b and 2e; Supplementary Figs. 10 and 11), suggesting that contemporary rice landraces retain genomic signals of past dispersal across Asia.

**Figure 2:**
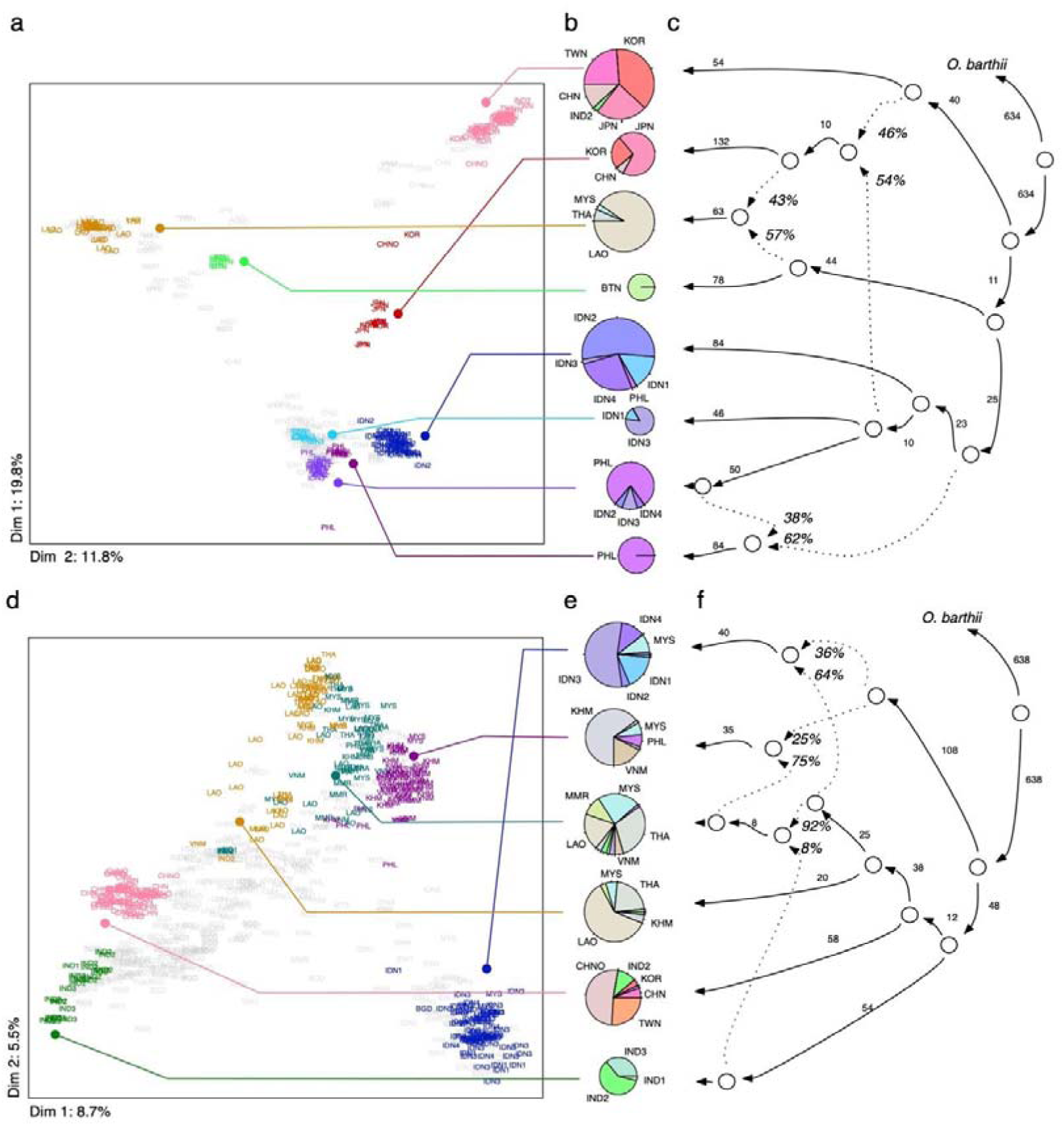
Japonica and indica rice subpopulations. (a) All japonica and (d) indica landraces projected onto first two dimensions after multidimensional scaling of genomic distances. (a) japonica genotypes were clustered using k-medoids (k = 9 subpopulations) and filtered using silhouette parameters, which resulted in K = 8 discrete subpopulations (colored labels). (d) indica genotypes were clustered using k-medoids (k = 7 subpopulations) and filtered resulting in K = 6 discrete subpopulations (colored labels). Pie charts representing the geographical composition of each discrete subpopulation of (b) japonica and (e) indica subgroups. Chart diameter is proportional to the number of individuals in each subpopulation. (c) Admixture graph for k = 9, K = 8 japonica subpopulations, rooted with *Oryza barthii* as an outgroup. This graph represents topology consistent between models for all lower k’s. (f) Best admixture graph for k = 7, K = 6 indica subpopulations, rooted with *O. barthii* as an outgroup. Although this represents the best model, it is not consistent with other topologies at lower k’s, likely due to complex history of indica. (c and f) Solid lines with arrowheads represent uniform ancestries (attached numbers show scaled drift parameter f_2_), while dashed lines represent mixed ancestries (% values indicate estimated proportion of ancestry).

## Relationships between japonica subpopulations

We modelled subpopulation relationships separately for japonica and indica using the admixture graph framework (*23*). We reconstructed relationships between japonica subpopulations at k = 2 to 9 considering graphs with population f-statistic z-scores <3. Throughout all k levels, we find two similar and consistent graph topologies (Fig. 2c; Supplementary Fig. 16), which we used to infer dispersal routes of japonica.

As expected (*2*), at k = 2 we observe divergence between lowland temperate varieties in Northeast Asia (Korea, Japan, China and Taiwan) and tropical varieties from the Malay Archipelago (Malaysia, Philippines and Indonesia). At k = 3, we find a major lineage of tropical upland japonica in mainland Southeast Asia as sister group to Malay Archipelago landraces or from admixture with an ancestral temperate lineage (Supplementary Figs. 10 and 16). At higher k, these mainland Southeast Asian upland landraces always incorporates admixture from an ancestral temperate japonica population (see below).

At k = 4 we observe separation of primarily Indonesian from Philippine and Bornean landraces. Subsequently, at k = 5, upland temperate japonica in Northeast Asia emerges as an admixture between lowland temperate and upland tropical varieties. Further increase of k allows separation of distinct Malay Archipelago subpopulations: a small subpopulation associated with the Philippines splits first, followed by a subpopulation in the Indonesian island of Java. Subsequent divisions among Malay Archipelago subpopulations are not fully resolved (Supplementary Fig. 16). Nevertheless, at k = 8, we identify a Bhutanese subpopulation closely related to upland Laotian landraces, and may represent a relict descendant population of the first early split in tropical japonica.

## The rise of temperate japonica

Combining genomic, geographic, archaeological and paleoenvironmental data, we reconstructed routes and timing of the ancient dispersal of rice in Asia. Japonica represents the first domesticated *O. sativa* (*9–11*), and its tropical form was cultivated in eastern China between the Yangtze and the Huang He (Yellow) river valleys (*13*). This occurred during the Holocene Climate Optimum (HCO), a period of increased monsoon activity and warmer temperatures between ∼9,000 and 4,000 yBP (*24, 25*); this coincides with the rise in frequency of non-shattering rice from ∼20% just after 8,000 yBP to fixation at ∼5,000 yBP (*5, 6*).

The first major population divergence in japonica separates temperate from tropical landraces (Supplementary Figs. 10 and 16). Using sequentially Markovian coalescent (SMC++), we estimated a cross-coalescence split time between temperate and tropical japonica at ∼5,000 to 1,500 years ago, with 75% of estimates between ∼4,100 to 2,500 years ago (Fig. 3a; Supplementary Fig. 17). Using dated archaeobotanical rice remains (*13*), we note that rice agriculture spread north- and eastward along the Huang He river (*26*) and westward into the Chengdu Plains and the Southwest China Highlands between ∼5,000 to 4,000 yBP (*27–29*)(Fig. 3b; Supplementary Fig. 18). During a minor climatic cooling event at ∼5,000 yBP, rice appears maladapted in parts of eastern China (*30*). In the Shandong Peninsula, rice disappeared by 5,000 yBP and briefly re-emerged 4,500 yBP as a short-grained variety similar to contemporary temperate japonicas (*31*). A global temperature decrease that followed the HCO at ∼4,200 years ago, the ‘4.2k event’ (*24, 25*), resulted in waning rice agriculture in East China and strong pressure for japonica to adapt to a temperate environment (*31*). Congruent with this, we observe that the highest density of estimated temperate japonica split times start at ∼4,100 years ago (Fig. 3a; Supplementary Fig. 17).

Temperate adaptation created opportunity for northeastern dispersal of japonica in Asia. From our demographic analysis of temperate japonica we note a ∼5-10-fold Ne reduction between ∼3,500 to 3,000 yBP (Fig. 3c; Supplementary Fig. 19), which we interpret as a founder bottleneck during expansion to its new temperate niche. Indeed, this is consistent with archaeological dates for the introduction of rice agriculture to Korea (*32, 33*) and Japan following decrease in rice remains in Eastern China (Supplementary Fig. 18).

**Figure 3:**
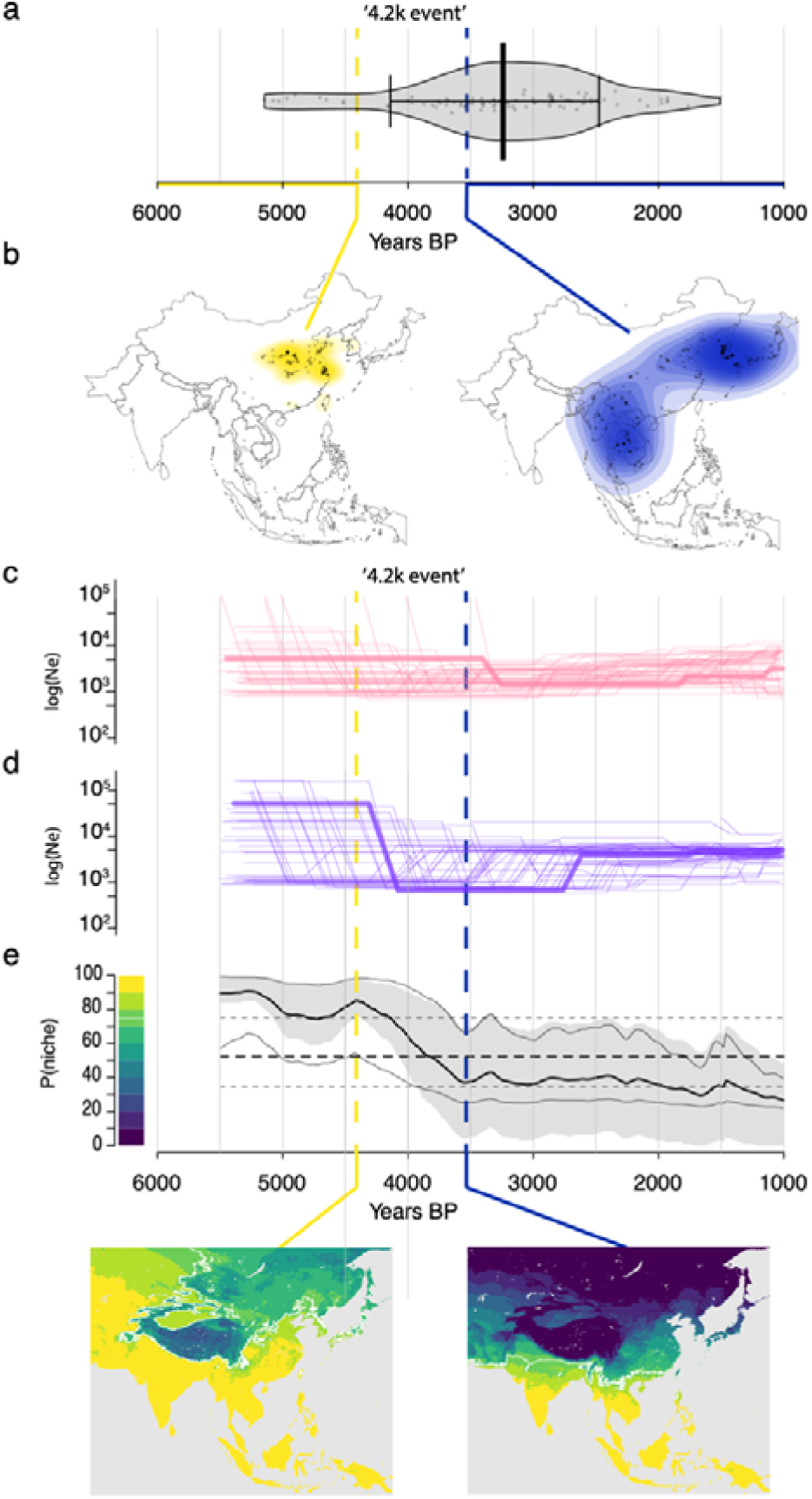
Demographic, paleoenvironmental and archaeological context of temperate japonica rice emergence. (a) The distribution of temperate-tropical split times estimated from cross-coalescence analysis carried out for 50 pairs of temperate and tropical individuals. (b) Maps indicating geographic locations and densities of archaeological sites with rice macro-remains. To the left: cumulative archaeobotanical evidence from 9,000-4,400 years BP, to the right: cumulative archaeobotanical evidence from 3,500-1,000 years BP. Effective population sizes over time in (c) tropical and (d) temperate japonica subpopulations. Thin lines represent demographic histories for 50 randomly sampled individuals, while bold lines represent joint models. (e) Probability of tropical rice being in the thermal niche (assuming requirement of 2900 growing degree days, at 10°C base) over time. The mean (thick black line) and the interquartile range, 25% to 75% (gray shaded area) of probability of being in the thermal niche. The thin black lines are the mean probabilities using the lower and upper confidence intervals of the temperature reconstruction. The two inset maps show the geographic distribution of niche probabilities; to the left: before climate cooling (4,400 years BP), to the right: after climate cooling (3,500 years BP).

## The southward spread of japonica

Throughout the HCO, tropical japonica was cultivated in eastern China; its contemporary descendants however, are grown predominantly in Southeast Asia (*1*), and we indeed find that Southeast Asian subpopulations descend from the tropical lineage that diverged from temperate japonica. Demography reconstruction at k = 2-4 shows that tropical japonica lineage experienced a ∼50-100-fold population (Ne) contraction between ∼4,500 to 4,000 yBP, and partial Ne recovery starting ∼2,500 yBP (Fig. 3d, Supplementary Fig. 19). The population contraction in tropical japonica is contemporaneous with the 4.2k event, raising the possibility that cooling explains the collapse of tropical rice cultivation in East Asia and its southern relocation. This coincides with the arrival of rice in the far south of China ∼4,500 yBP and a shift to rainfed, upland cultivation (*34*).

Given the importance of temperature in shaping japonica genomic diversity across its geographic distribution (Fig. 1g), we used a thermal niche model (*35*) based on reconstruction of Holocene temperatures (*36*) to estimate the probability of tropical rice cultivation in different areas during the post-HCO period (Fig. 3e; Supplementary Fig. 20). Survival probabilities of tropical japonica between ∼4,400 and 3,500 yBP dropped dramatically in eastern China and high-altitude South China (survival probability < 50%) compared to Southeast Asia [survival probability > 90%](Fig. 3e; Supplementary Video 1). Indeed, after the cooling period we observe high densities of archaeological rice remains in Southeast Asia (Fig. 3b; Supplementary Fig. 18).

After the HCO, rice dispersed from China to Southeast Asia into Laos and Bhutan, and through maritime routes to the Philippines, Malaysia and Indonesia. In our admixture graph analysis, we find an early split in the tropical lineage that separates Bhutan and Laos upland rice from rice in the Malay Archipelago (Fig. 2c). From coalescence analyses we observe a ∼50-100-fold population contraction in the remote upland (Bhutan) rice population between ∼4,000 and 3,000 yBP (Fig. 4; Supplementary Fig. 19), which may arise from a bottleneck associated with population movements into these new areas. Emergence of upland rice in Laos and Bhutan coincides in time and space with widespread establishment of rainfed rice agriculture in mainland Southeast Asia, ∼4,000 yBP (*12, 37*) and dispersal of metallurgy traditions from Bronze Age Yunnan, ∼3,500 yBP southwards to Thailand by ∼3,000 yBP (*38, 39*). Subsequent agricultural intensification of rice production took place from ∼2,500 to 1,500 yBP and included evolution of irrigation systems in present-day Thailand (*40*). Consistent with these, ancient human DNA studies in Southeast Asia report two farmer-associated migration events from East Asia, one at least 4,000 years ago and a second before 2,000 yBP (*41, 42*).

**Figure 4:**
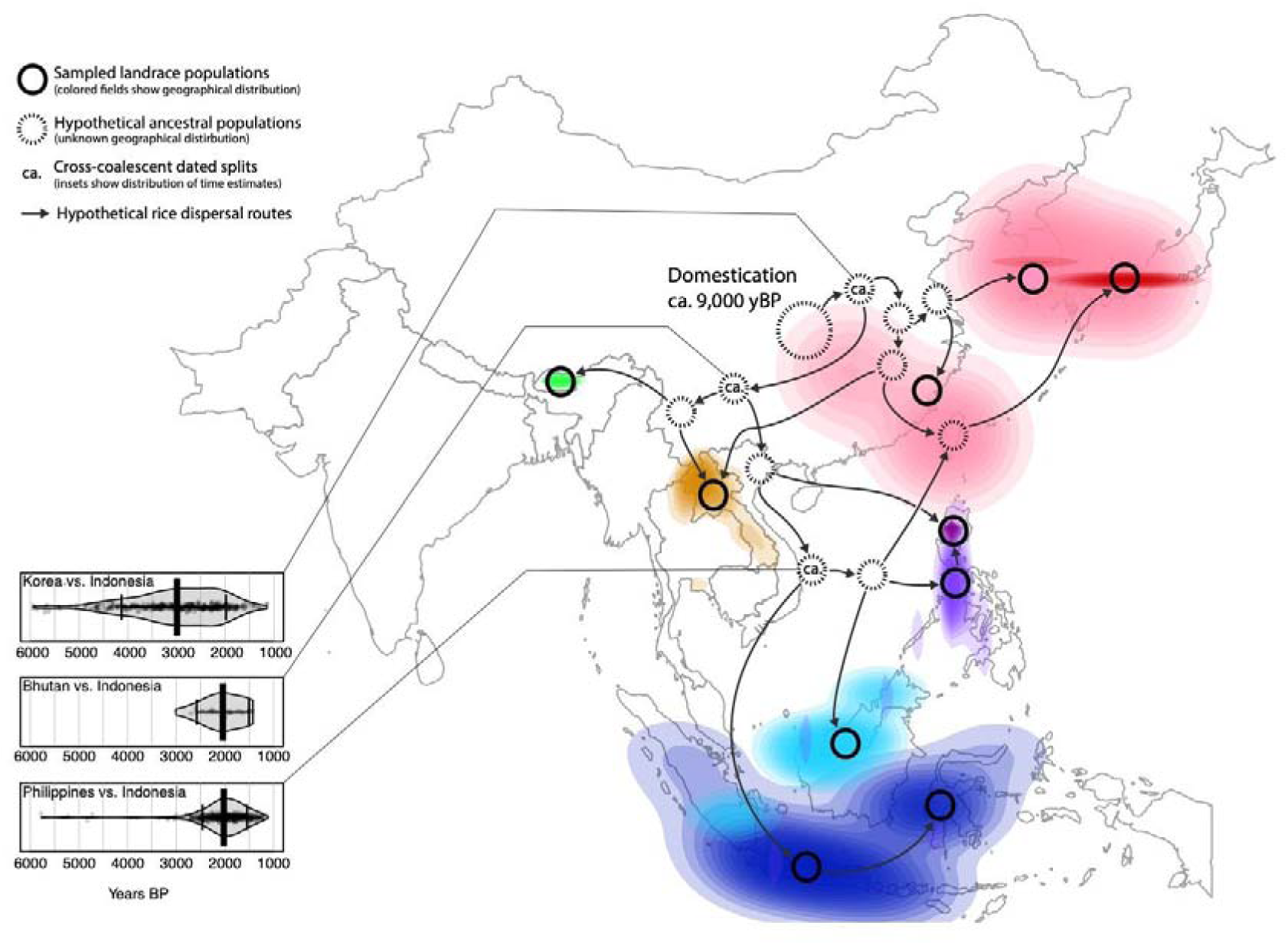
Proposed dispersal map of japonica rice in Asia. Map generated for japonica, K = 8 discrete subpopulations. The geographic distributions of subpopulations were represented as colored, two-dimensional Kernel density fields. Bold circles represent leaves in the admixture graphs and are mapped close to the centers of subpopulation distributions. Dashed circles represent hypothetical ancestral subpopulations inferred from splits in best-matching admixture graphs; their precise geographic placement is uncertain. The distribution of split times between non-admixed subpopulations was created from cross-coalescence estimates summarized over all k levels and presented as violin plots. Arrows indicate hypothetical routes of dispersal.

Our analysis also shows an ∼5-10-fold Ne decrease in the Malay archipelago between ∼3,000 and 2,500 yBP, and based on cross-coalescence analyses, divergence between mainland and Malay Archipelago rice occurred between ∼3,000 to 1,500 years ago (75% of estimates in ∼2,500 to 1,600 yBP) [Fig. 4; Supplementary Fig. 19]. Distinct island populations in the Malay Archipelago diverged at around a similar timeframe, in an interval from ∼3,000 to 1,000 years ago (75% estimates fall between ∼2,500 and 1,500 yBP). This period coincides with dispersal of Dong Son drums in the Malay Archipelago (∼2,400 years ago) (*38, 43*), and suggests maritime dispersal of rice from a North Vietnam hub within the Austronesian Trading Sphere, which stretched between Taiwan and the Malay Peninsula (*44, 45*). Ancient DNA studies also suggest a wave of Austronesian human expansion into island Southeast Asia ∼2,000 years ago (*41*), which agrees with our estimates of japonica movement into the area. Interestingly, upland temperate japonica in Japan appears to be an admixed population of local lowland temperate rice and upland tropical rice from the Malay Archipelago which may have moved northwards through Taiwan and perhaps the Ryukyu Islands ∼1,200 yBP (*46*).

## Relationships and dispersal of indica subpopulations

We reconstructed relationships between indica subpopulations with k = 2 to 7. Divergence between Sino-Indian and Southeast Asian indica is present in all graph topologies beginning at k = 2 (Supplementary Fig. 21). At k = 3 we observe separation of mainland and island Southeast Asian subpopulations, while at k = 4 we observe separation of Indian from Chinese landraces. With k = 5 and k = 7 we note differentiation of mainland Southeast Asian landraces into subpopulations associated with Laos, Thailand and Cambodia (Fig. 2f). Interestingly, a subpopulation associated primarily with Cambodia, and another in Indonesia, share ancestry with the main Laos/Thailand Southeast Asian lineage as well as an early ancestral indica population. Increasing to k = 8 also increases the number of admixture events in the model to four, which renders further exhaustive graph topology searches unfeasible.

Higher diversity of graph topologies in indica, likely due to weaker population structure and elevated gene flow (Supplementary Figs. 14 and 15), makes it difficult to reconstruct indica dispersal routes. Moreover, given the complexity in multiple reconstructed admixture graph topologies, we can only confidently date separation of Chinese and Indian indica, which is unaffected by admixture. Our analysis estimates this divergence at ∼2,500 and 1,100 yBP (75% of estimates between ∼2,000 and 1,400 yBP)[Fig. 5; Supplementary Fig. 17]. Possible routes for indica dispersal from India to China could be the Silk Road or more direct passage to Southwest China across the Hengduan mountains. The timing agrees with written reports of the introduction of Buddhism from India to China at ∼1,950 yBP (*47*), but is later than the earliest putative finds of indica rice in China (*48*). The close relationship between Indian and Chinese subpopulations is mirrored by higher proportions of irrigated varieties in both regions; in contrast, Southeast Asian varieties are more often rainfed (*19*).

**Figure 5:**
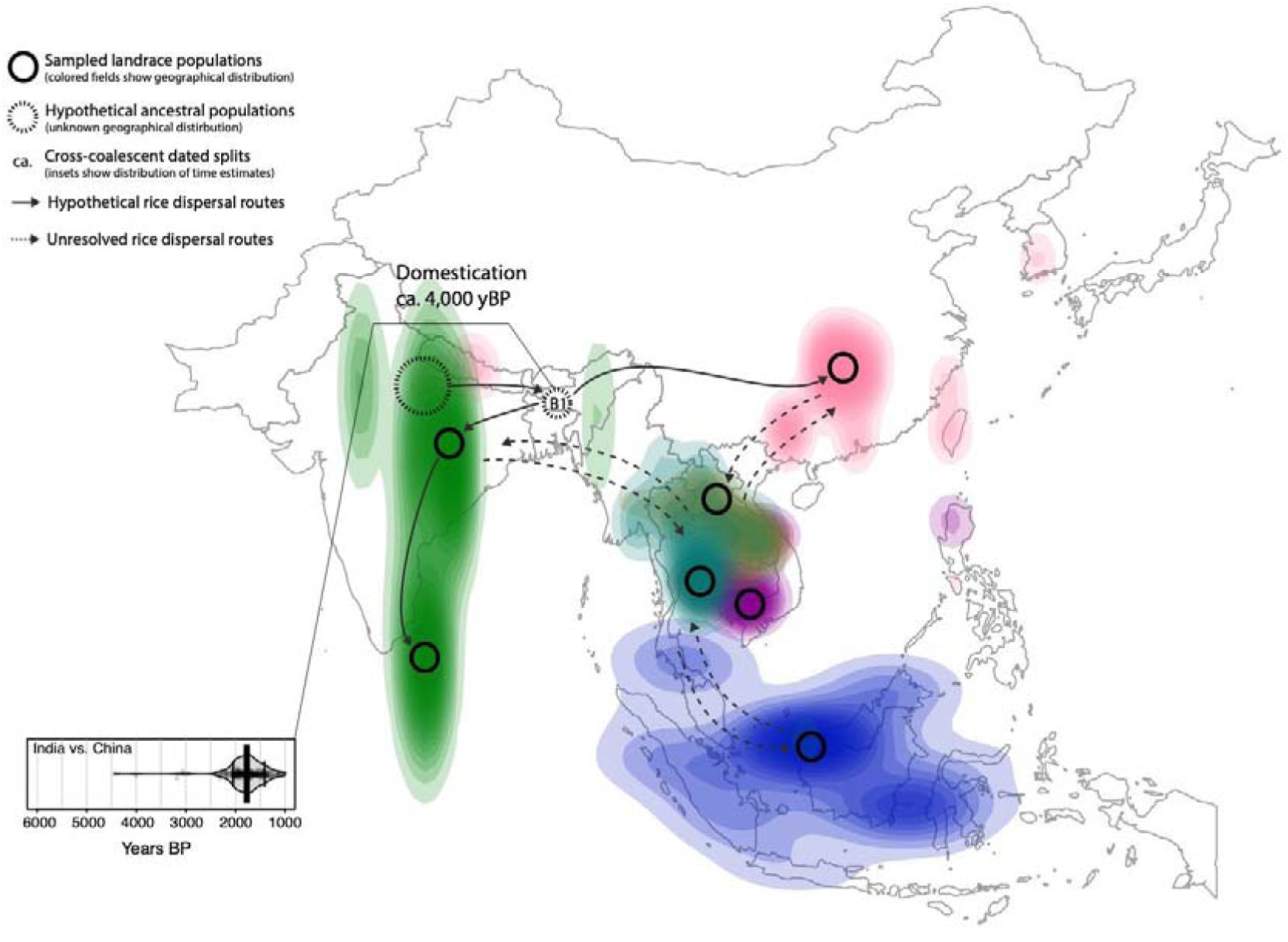
Proposed dispersal map of indica rice in Asia. Map generated for indica, K = 6 discrete subpopulations. The geographic distributions of subpopulations were represented as colored, two-dimensional Kernel density fields. Bold circles represent leaves in the admixture graphs and are mapped close to the centers of subpopulation distributions. Dashed circle represents consistent split; its geographic position is uncertain. The distribution of split times between non-admixed subpopulations was created from cross-coalescence estimates summarized over all k levels and presented as violin plots. Solid arrows indicate hypothetical routes of dispersal, while dotted arrows indicate possible routes that remain unresolved from admixture graphs.

Indica dispersal to Southeast Asia (e.g., Thailand and Cambodia) were either from India or China (Fig.5; Supplementary Fig. 22). From archaeobotanical studies, indica arrived in Central Thailand at ∼1,800 years ago (*40*), at a time when Asian trade routes were well established (*12*). Late adoption of indica in Southeast Asia is hypothesized to be due to early availability of japonica in this region (*12*). There is no earlier archaeological evidence for indica cultivation in Southeast Asia, and hence it comes as a surprise that indica mainland subpopulations suffered dramatic population size reduction between ∼5,000 and 3,500 yBP (Supplementary Fig. 23). It is even more puzzling that a bottleneck in indica subpopulation in Indonesia occurred between ∼6,000 and 5,000 yBP, suggesting complex origins, perhaps partially from local wild ancestors or managed pre-domesticated varieties (Supplementary Fig. 22).

## Summary

Rice domestication in the Yangtze Valley had an enormous impact on the peoples of East, Southeast and South Asia. In the first ∼4,000 years of its history, Japonica rice cultivation was largely confined to China, and its dispersal and diversification did not occur until the global 4.2k cooling event. This abrupt climate change event, which was characterized by a global reduction in humidity and temperature, had widespread consequences: it is believed to have caused the breakdown of rice agriculture in East Asia (*24, 31*), turnover of cattle ancestry in the Near East (*49*), and the collapse of civilizations from Mesopotamia (*50*) to China (*51*). We find from our genomic and paleoclimate modelling that the 4.2 k event coincides with the rise of temperate japonica and the dispersal of rice agriculture southwards into Southeast Asia. Moreover, indica began to be domesticated in South Asia at around this period, and spread later into China and Southeast Asia. Correlation between changing climate and rice distribution raises the possibility for a causal relationship, and indeed we find temperature is a key environmental factor patterning contemporary rice genomic diversity.

The ability to infer dispersal patterns of rice arises from the availability of extensive landrace populations, whole genome sequences and population genomic approaches, as well as environmental, archaeobotanical and paleoclimate data. Reconstructing the history of domesticated species provides insight into the evolutionary process, nature of human/plant co-evolutionary dynamics, and extrinsic landscape, environmental, and cultural factors that drive crop dispersal. Armed with knowledge of the pattern of rice dispersal and environmental features that influenced this migration, it may be possible to examine the evolutionary adaptations of rice as it spread to new environments, which could allow us to identify traits and genes to help future breeding efforts.

## Acknowledgments

We would like to thank our colleagues for helpful discussions on this project.

## Author contributions

RMG and MDP conceived and designed the study with input from JRL and SCG. JYC, ISP and OW generated sequencing data. MDP, SN and MMO supervised laboratory work. RMG assembled and processed the sequencing data. SCG and ESB assembled and processed the environmental data with input from JRL. JRL lead the spatial analyses with input from RMG. ESB and ERS carried out travel time analyses with input from JRL. JRL carried out RDA analyses. RMG carried out population structure, admixture graph and coalescence analyses. RKB and JAdG conducted thermal niche modelling. DQF, CCC and JAdG provided archaeological context. MDP, RMG and JRL wrote the manuscript with input from all authors.

## Data and materials availability

Raw FASTQ reads for 178 accessions whose genomes were re-sequenced for this study have been deposited in the Sequence Read Archive under SRA Bioproject accession numbers PRJNA422249 and PRJNA557122. Sources for all downloaded data are available in the supplementary materials. Code repositories are refered in the supplementary materials

