## Supplemental information for "Genomic history and ecology of the geographic spread of rice"

^1^Center for Genomics and Systems Biology, New York University, New York, NY 10003 USA; ^2^Department of Biology, Pennsylvania State University, University Park, PA 16802 USA; ^3^Instituto de Tecnologia Química e Biológica António Xavier, Universidade Nova de Lisboa, Av. da República, 2780-157 Oeiras, Portugal; ^4^Crow Canyon Archaeological Center, Cortez, CO 81321, USA; ^5^Carnegie Mellon University Libraries, Pittsburgh, PA 15213-3890; ^6^Department of Biological Sciences, University of Manitoba, Winnipeg, MB R3T 2N2, Canada; ^7^Institute of Archaeology, University College London, London, WC1H 0PY, United Kingdom; ^8^School of Cultural Heritage, North-West University, Xi’an, Shanxi 710069, China; ^9^School of Biology and Environmental Science, University College Dublin, Belfield, Dublin 4, Ireland; ^10^Department of Anthropology and Scripps Institution of Oceanography, University of California San Diego, La Jolla, CA 92093, USA; ^11^Center for Genomics and Systems Biology, New York University Abu Dhabi, Saadiyat Island, Abu Dhabi, United Arab Emirates. ^12^Institute for the Study of the Ancient World, New York University, New York, NY 10028.

*Corresponding author

**Keywords**: domestication, crop evolution, diversification, paleoclimate, archaeobotany, admixture analysis, coalescence, *Oryza sativa*

TABLE OF CONTENTS

**1. Materials and methods	2**

**1.1. Data collection	2**

*Landrace status	2*

*Geolocations and cultivation systems	2*

*Biotic variables	3*

*Abiotic variables	4*

*Sequencing data	5*

**1.2. Processing of sequencing data	6**

*Alignment and genotyping	6*

*SNP filtering	6*

**1.3. Genomic variance partitioning	8**

*Landrace subsetting	8*

*Migration barriers	8*

*Spatial correlations	9*

*Redundancy analyses	10*

**1.4. Discrete population structuring and dispersal	12**

*Clustering and discretization	12*

*Admixture graph reconstruction	13*

*Demography and split time reconstruction	14*

**1.5. Archaeological and paleoenvironmental context	15**

**2. Supplementary references	17**

**3. Supplementary figures	22**

**1. Materials and Methods**

**1.1. Data collection**

*Landrace status*

We considered 2,466 domesticated Asian rice (*Oryza sativa* L.) accessions from the International Rice Genebank Collection (IRGC) at the International Rice Research Institute (IRRI) that were included in the 3K-RG project (*1*), as well as an additional 178 accessions that were re-sequenced at New York University (Supplementary Table 1).

The definitions of landrace are very complex (*52*, *53*) and hard to apply in practice during material collections. In our work we relied on the fact that landraces contain the signal of association with local geographic, environmental and cultural context. To that end we used the following criteria: 1) Pre-selected ‘candidate’ landraces from available annotation, and 2) filtered them based on their joint genetic and geographic clustering.

Accession passport data were obtained from the International Rice Information System (<http://iris.irri.org/>) (*54*). We considered sample status of each accession and removed ‘improved variety’, ‘wild’ and ‘weedy’ accessions, while keeping ‘traditional variety/landrace’ accessions. We also kept ‘breeding/inbred line’ accessions if these were pure lines directly derived from ‘traditional varieties/landraces’ or were classic breeding lines from before the Green Revolution in the 1960s. From that set we removed any genetic clusters that were represented by individuals collected in in countries that do not share borders.

*Geolocations and cultivation systems*

Landrace geo-references were obtained Genesys (<https://www.genesys-pgr.org/welcome>). For some landraces, instead of precise geo-coordinates, country- or region-level centroids were given (Supplementary Table 1). This problem was particularly relevant for landraces from China and Japan. Data on agro-ecosystems in which accessions are cultivated were obtained from IRIS (*54*). Based on their agro-ecosystem of origin, accessions are divided into six cultivation types: ‘irrigated’, ‘rainfed lowland’, ‘deepwater’, ‘upland’, ‘tidal wetland’, and ‘swamp’ (*55*).

Accession growing season(s) in its local environment were estimated through considering information on cultivation type with prevalent rice growing season months at the collection location. The latter information was obtained from the Rice Almanac (*56*), and Rice Atlas (*57*). An accession from ‘irrigated’ agro-ecosystem was assumed to be grown in all growing seasons if there were multiple seasons in its location of origin, since sufficient irrigation can presumably be provided in ‘off’ or ‘dry’ seasons. Accession from other agro-ecosystems were assumed grown only in the ‘main’ or ‘wet’ growing season as indicated in the Rice Almanac (*56*), and the Rice Atlas (*57*). An accession’s growing season months were further specified if additional metadata on growing season was available from the IRIS database (*54*).

*Biotic variables*

It has been suggested that wild relatives of rice, particularly *Oryza rufipogon* and *O. nivara* hybridized with cultivated rice in the past (*9*–*11*) altering the genomic composition of local subpopulations. We therefore considered the wild gene pool available to each candidate landrace. To that end we used published ancestry composition data for rice wild relatives (*58*). For each of our candidate landraces we took the ten geographically closest wild individuals and calculated their mean ancestry composition.

We also considered rice-human relationships not covered by geographic distance/resistance dispersal. One important rice grain property in a culinary cultural context is stickiness, which is determined by the *waxy* gene (*59*). As a proxy for conscious cultural preferences, we genotyped *waxy* alleles from genome-wide data. We also considered effects of unconscious cultural preferences on distribution of rice genomic diversity by accounting for the language family of nearby human populations. To this end we downloaded the linguistic map from the Glottolog database (*60*) and for each candidate landrace we queried the geographically closest spoken language.

*Abiotic variables*

We collated data for a suite of climate-related variables at the geo-location of each landrace using the EXTRACT function of the R package RASTER (*61*) v.2.8-19. Six temperature variables (average coldest temperature throughout growing season(s), average coldest temperature in first two months of growing season(s), mean temperature over growing season(s), average high temperature for last two months of growing season(s), growing degree days [GDD] in growing season(s), and inter-annual coefficient of variation of GDD) and three precipitation variables (accumulated precipitation in two months before the growing season(s), mean precipitation throughout growing season(s), and inter-annual coefficient of variation of precipitation) were derived from CHELSA climatological data (v1.2), which provides monthly and mean annual precipitation and temperature data at 30 arc-second resolution for the time period 1979 to 2013 (*62*). For calculations of GDD, we used monthly means as proxies of average daily air temperatures for months in the growing season with a mean above a base temperature of 10°C.

We included two variables that reflect evapotranspiration processes during the growing season(s): potential evapotranspiration [PET] and ratio of PET to mean precipitation. PET variables were based on monthly values from the CGIAR-CSI Global-PET Database (*63*, *64*), <http://www.cgiar-csi.org>).

We also included distance from the geo-location of each landrace to the nearest lake or river based on a previous global analysis of human population distance to freshwater (*65*). Elevation above sea level was obtained from WorldClim (*66*).

Among edaphic variables, we included: soil salinity (measured as electric conductivity), pH, and sodicity (exchangeable sodium percentage) from the Harmonized World Soil Database v1.2 (<http://webarchive.iiasa.ac.at/Research/LUC/External-World-soil-database/HTML/index.html?sb=1>). Information on soil total nitrogen density was extracted from the Global Gridded Surfaces of Selected Soil Characteristics dataset (*67*). To capture the soil moisture potentially available for plant growth, we used plant extractable water capacity of soil (*68*) and depth to water table (*69*).

*Sequencing data*

Sequencing data for individuals that were marked as candidate landraces (see section 1) from the 3K-RG project were downloaded in fastq format from the Short Read Archive (SRA) using FASTQ-DUMP tool with option to split reads into forward, reverse and trimmed.

We generated sequencing data for additional 178 landraces. Leaf samples were ground using mortar and pestle in liquid nitrogen. DNA was extracted using the Qiagen DNeasy Plant Mini Kit following the manufacturer’s protocol (QIAGEN, Hilden, Germany). Yields ranged between 3 ng/ul and 102 ng/ul. Extracted DNA from each sample was prepared for Illumina genome sequencing using the Illumina Nextera DNA Library Preparation Kit. Sequencing was done on the Illumina HiSeq 2500 – HighOutput Mode v3 with 2×100 bp read configuration, at the New York University Genomics Core Facility. Sequencing data these accessions are available from the SRA under Bioproject accession numbers PRJNA422249 and PRJNA557122.

**1.2. Processing of sequencing data**

*Alignment and genotyping*

We used Nextflow (*70*) to build a pipeline for calling SNPs in our dataset (<https://github.com/grafau/NextGatkSNPs>). All steps necessary to obtain our SNP set are described below. Sequencing data in fastq format for each run of candidate landraces were mapped against the reference genome of indica variety Shuhui498 v.1.0 (*71*) using the global aligner BWA v.0.7.15 in ‘mem’ mode (*72*) and sorted using PICARD v.2.15.0. Sequences for the same sample, but from different runs, were merged and amplification duplicates were removed using PICARD. The resultant sam format files were validated and indexed producing bam format files.

Bam files were used to call haplotypes in GATK v.3.8 (*73*) with the HAPLOTYPECALLER function in ‘discovery’ mode and set to produce gvcf format files. Subsequently, gvcf files were validated and combined into eight batches with GATK, each batch containing approximately 200 landraces. These combined gvcf files were compressed and indexed using BGZIP and TABIX, respectively (*74*). Contents of combined gvcf files were divided into 12 chromosomes and each chromosome file was genotyped for all eight batches together using GATK’s GENOTYPEGVCFS function to produce the raw set of SNPs segregating among rice landraces.

*SNP filtering*

The raw set of SNPs was subject to a series of filtering steps. First, we only kept biallelic SNPs. Subsequently, we applied five filtering criteria: qualities normalized by depth (QD), mapping quality (MQ and MQRankSum), read position bias from Wilcoxon’s test (ReadPosRankSum), and strand bias from Fisher’s test (FS). Filtering thresholds for these criteria were trained dynamically using GATK’s VARIANTRECALIBRATOR function referencing a true-positive set of SNPs that were discovered independently in the 3K-RG project (*1*), and in the rice diversity panel (RDP) that was genotyped with a high-density SNP array (*75*). We applied the dynamic filter to our raw set of SNPs using GATK’s APPLYRECALIBRATION function conservatively set to recover 90% of true positives.

To obtain an estimate for expected heterozygosity in rice populations we calculated inbreeding coefficients in all landraces of *circum*-aus, indica, and japonica groups. Coefficients were calculated as medians of ratios, where each ratio equals observed heterozygosity divided by expected heterozygosity for each SNP with >5% minor allele frequency (only ratios smaller than 1 were taken into account). We then compared observed heterozygosity to expected heterozygosity for each SNP, given the inbreeding coefficient, and carried out a chi-square test to filter out SNPs with excess heterozygosity. We performed this step for all landraces and for each subgroup separately. We interpret excessively heterozygous sites as mis-mapped reads in chromosomal regions with structural variants that are present in the re-sequencing data but absent in the reference genome.

Next, we transformed vcf files into bed format files using PLINK v.1.90b4 (*76*, *77*), and kept only candidate landraces that were collected in Asia. From this set, we filtered out any SNP that had a lower than 80% genotyping rate with PLINK. This step was carried out independently for all landraces, and for indica and japonica subgroups separately. For some analyses (Supplementary Fig. 1) SNP sets were subject to additional two-step linkage disequilibrium pruning. The first step was carried out with the ‘INDEP-PAIRWISE’ function in windows of 10 kb with variant shift = 1 and r^2^ = 0.8. The second step was carried out with the same function in windows of 50 variants.

**1.3. Genomic variance partitioning**

*Landrace subsetting*

We used metadata retrieved from the online platform Genesys (for details see section 1) to annotate each candidate landrace with country of origin and subgroup designation (indica, japonica, *circum*-aus, *circum*-basmati) provided by the 3K-RG and RDP (*1*, *75*) projects. We then carried out factorial analysis of molecular variance in R v.3.4.1 (*78*) using the ADONIS function from the VEGAN package (*79*). Subsequently, we split the dataset into four subsets corresponding to four subgroups (each SNP set was subject to heterozygosity and genotyping rate filtering, see section 2). We used pairwise genomic distances among all landraces to calculate silhouette scores (*21*) for each landrace given its subgroup affiliation. We then filtered out landraces with silhouette scores below 0.2, as this might indicate admixture between subgroups or mislabelling. All analyses described in this section were carried out after phasing imputation on the SNP sets with BEAGLE v.5.0 (*80*).

*Migration barriers*

We estimated effective migration surfaces using the EEMS tool (*15*). We chose map outline coordinates that stretch from Pakistan to Japan and Papua New Guinea using an online tool (<http://www.birdtheme.org/useful/v3tool.html>) and specified a triangular grid with 200 demes for Voronoi tessellation. The best-fitting model was acquired from converging three independent runs of 5 million Monte-Carlo Markov Chain iterations of which the first 2 million burn-in runs were discarded. Surfaces were plotted in R v.3.4.1 (*78*) using the EEMS.PLOT function from the REEMSPLOTS package (*15*) and mapped with Mercator projection.

Fastest travel time between each pair of geo-referenced accessions was estimated using least-cost paths analysis in R v3.5.1 with the package GDISTANCEv1.1 (*81*). Traveling speed over land given the slope between adjacent grid cells was calculated according to Tobler’s Hiking Function (*82*), based on elevation data at 30 arc second resolution from WorldClim v1.4. For travel over sea, we assumed a constant speed of 3 knots under sail (*16*, *83*, *84*). GPS coordinates for each landrace accession were rounded to the nearest 0.1 degree to reduce computation time needed. Pairwise resistance distance matrices were populated separately for indica and japonica accessions. For details see: <https://github.com/em-bellis/riceTravelTime>

*Spatial correlations*

We tested whether genetic distance between landraces could be explained better by geographic distance or estimated travel time between geo-locations of origin. We first filtered out landraces in China, because they all were annotated with low resolution geo-coordinates that mapped to country and regional centroids. We employed linear mixed model with maximum likelihood estimation of Clarke *et al*. (*85*) and used Akaike Information Criterion (AIC) (*86*) to select between geographic distance versus travel time models. This linear mixed model includes spatial random effects to account for non-independence among nearby samples. The use of AIC with such a mixed model has been shown to offer the greatest accuracy in identifying the true isolation model under a wide range of scenarios (*87*). We implemented our mixed model and AIC calculations with RESISTANCEGA package (*88*, *89*) in ‘R’ v.3.6.0 (*78*). Proportions of variance explained (r^2^) were calculated with the LM function and p-values were calculated using Mantel tests and permutations implemented in VEGAN (*79*).

Processes driving gene flow may have been very different in continental Asia versus the Malay Archipelago. Additionally, the travel time model we developed was new, and therefore had an uncertain ability to capture different travel mechanisms. Thus, we stratified analyses into two main groups each for both japonica and indica: a group of ‘continental’ and a group of ‘island’ landraces. Continental landraces were defined to include those north of 9.7ºN latitude and west of 110ºE longitude, thus excluding the relatively small number of isolated mainland landraces to the east (e.g. eastern China). Island landraces included those from the Malay Archipelago and the Malay Peninsula, but not from the islands to the north (*i.e.*, Taiwan or Japan).

*Redundancy analyses*

Redundancy analyses (RDA) are eigenanalyses for multivariate responses and multivariate predictors that maximize the proportion of variation explained in the responses. We used RDA to identify sets of variables important for explaining SNP variation in landraces and for identifying specific (a)biotic variables explaining the most genome-wide SNP variation. To incorporate pairwise geographic distance or travel time into our RDA, we converted distance matrices into spatial weighting matrices and then a reduced-dimension set of orthogonal variables (Moran’s, eigenvector maps, MEMs)(*90*). MEMs are eigenvectors of the pairwise spatial weighting matrix among samples. We optimized both geographic distance and travel time matrices using a subset of 10,000 randomly chosen SNPs for response variables in RDA, optimizing separately for japonica and indica.

Weighting matrices among unique landrace collection locations (Chinese accessions were all filtered out) were generated using ADESPATIAL package (*91*) in R. We used two algorithms, Gabriel graph and distance-based graph, to generate three candidate connectivity matrices. The Gabriel graph results primarily in connections among neighboring sites. A distance-based graph connects sites closer than a given threshold, for which we used two values: minimum distance required to connect all points (*i.e*., the largest distance of a minimum spanning tree) and infinity (resulting in a fully connected graph (*90*)). With each of these three connectivity matrices we generated two spatial weighting matrices using two distance decay functions: linear (weight between two sites = 1 - *D*/*D*_max_ where *D* is distance between sites and *D*_max_ is maximum distance among all sites) or concave up (weight between two sites = *D*^-0.01^). These connectivity and weighting algorithms resulted in six diverse MEM sets, differing largely in levels of spatial autocorrelation and structure among MEM eigenvectors. We used the Bauman *et al*. (*90*) forward-selection of MEM eigenvectors algorithm to optimize number of eigenvectors (restricted to those with positive eigenvalues) included in RDA for each MEM set. Optimization is based on adjusted r^2^ (which are penalized/adjusted for number of explanatory values), and the MEM set with greatest adjusted r^2^ is defined as the optimal set. In the RDA presented in the main text we used weighting matrices based on geographic distance for indica and travel time for japonica, because model selection favored these distance measures. For indica and geographic distance, optimization selected 25 MEM eigenvectors from the connectivity matrix based on connecting all sites within the threshold distance required to connect all points in a single graph and using weighting that was a linear function of distance. For japonica and travel time, optimization selected the same connectivity matrix and distance weighting algorithms, with 33 eigenvectors. These eigenvectors were included in the RDA described below on japonica and indica whole SNP dataset.

We then conducted RDA with variance partitioning (*17*) to quantify proportion of genome-wide SNP variation explained by each of four categories of covariates: abiotic variables, geographic isolation MEMs, *waxy* allelic status, and language family. Variance partitioning estimates proportion of SNP variance explained by variables in each category and by collinearity among variables. To identify specific abiotic variables associated with genome-wide divergence among landraces, we also conducted RDA using only abiotic gradients for indica and japonica. For visualization, specific abiotic variables highlighted in Fig. 1 in the main text indicate those loading most strongly in each direction along each RDA canonical axis as well as those loading most strongly in each diagonal (identified by multiplying the loadings on the first two canonical axes). All RDA (including variance partitioning) were conducted using VEGAN (*79*).

**1.4. Discrete population structuring and dispersal**

*Clustering and discretization*

Clustering was visualized using multidimensional scaling methods. Genetic distances among and within each rice subgroup were calculated between all pairs of candidate landraces using PLINK v.1.9 (*76*, *77*) with formulation: 1 - IBS, where IBS is identity-by-state. After importing the distance matrix into R v.3.4.1 (*78*) the CMDSCALE function was used to calculate eigenvectors (*92*), which were plotted in three dimensions. The variance explained by each dimension was calculated as the dimension’s eigenvalue divided by sum of all positive eigenvalues.

Formal clustering of landraces within japonica and indica was carried out based on pairwise genetic matrices with the partitioning around medoids (PAM) method (*93*) implemented as the PAM function in CLUSTER package for R v.3.4.1 (*78*). Subsequently, clusters were filtered with our DISCRETIZE algorithm implemented in R. The algorithm first removes individuals with negative silhouette scores. Second, for each cluster it designates a pairing partner, which is another cluster with the least-distant medoid. DISCRETIZE simulates individuals that are admixed between the two paired clusters with requested ancestry proportions by computing weighted-mean distance between paired medoids and all other individuals (here we simulated individuals with 0.5-0.5, 0.4-0.6, and 0.6-0.4 admixture proportions). For all simulated individuals, our algorithm computes silhouette scores and keeps the highest value as threshold for filtering. Individuals are clustered with PAM and filtered based on each cluster’s silhouette threshold. This process is repeated iteratively until no more individuals are filtered out. A script written in ‘R’ that can perform these analyses is publicly available (<https://github.com/grafau/discretize>).

Clustering and discretization was carried out independently for a number of clusters, k, that varied from 2 to 12. Discrete clusters are considered subpopulations and their members are considered landraces conditional on a co-localized geographic distribution within each discrete cluster. We investigated composition of clusters with regard to region of origin to see if each fulfilled our latter criterion for landrace status. One indica cluster exhibited poor geographic co-localization and was therefore removed from all analyses (see section 1). In order to visualize the geographic provenance of each discrete cluster, we plotted the two-dimensional distribution (for latitude and longitude) of landraces using the GEOM_DENSITY2D and STAT_DENSITY2D functions from the GGPLOT package (*94*) onto a map of Asia in R v.3.4.1 (*78*).

*Admixture graph reconstruction*

We reconstructed admixture graphs for japonica and indica subpopulations defined by the DISCRETIZE algorithm. Reconstruction attempts were carried out independently for varying numbers of subpopulations, with k ranging from 2 to 9, using 19 accessions of *Oryza barthii* as outgroup. The CONVERTF function from ADMIXTOOLS was used to produce eigenstrat data files, and QPGRAPH function was used to evaluate whether models fit the data. Models were taken from ADMIXTUREGRAPH package (*95*) in R v.3.4.1 (*78*) and transcribed into the format accepted by ADMIXTOOLS (*23*).

From k = 2 to k = 5 (3 to 6 subpopulations including outgroup) we explored the entire space of possible models with 0, 1, and 2 migrations and reported all models with f_4_-statistic z-scores < 3.0 (Supplementary Fig. 15 and 16). For higher k’s we explored all possible models with 6 subpopulations and 0, 1, and 2 migrations, keeping only those with f_4_-statistic z-scores < 3.0. For each model we kept, we attached an additional subpopulation in all possible nodes using ADMIXTUREGRAPH and tested the resultant models in ADMIXTOOLS, again keeping only models with f_4_-statistic z-scores < 3.0. We progressively added subpopulations until no more were present or until no models with f_4_-statistic z-scores < 3.0 were found. In the latter case, we kept all models with f_4_-statistic z-scores lower than 10.0. We then added an additional admixture event in all possible nodes using ADMIXTUREGRAPH and tested resultant models in ADMIXTOOLS, keeping only models with f_4_-statistic z-scores < 3.0. The number of possible models fulfilling this criterion was large, so we summarized their topologies in three different ‘topology groups’ and showed representative models characterized by the best z-scores together with total number of models in these topology groups (Supplementary Fig. 15 and 16).

*Demography and split time reconstruction*

To better understand past demographics of rice, we attempted to reconstruct past effective population sizes using the Sequential Markov Coalescent method (SMC++) (*96*). Reconstructions were carried out independently for a varying number of subpopulations, with k ranging from 2 to 9. We selected a variety of ‘distinguished pairs’ for each subpopulation through sampling 50 individuals without replacement and pairing them with 50 individuals sampled with replacement. In subpopulations with fewer than 50 individuals assigned, we sampled all of them and paired each with individuals sampled with replacement. We then partitioned vcf files into smc haploblock files for each distinguished pair, further partitioned for each chromosome, and masked the homozygous pericentromeric regions (*97*). Subsequently, we used a polarization error of 0.5 and mutation rate of 6.5 x 10^-9^ (*98*) in the ESTIMATE function of SMC++ to estimate past effective population sizes. Results were scaled in time using an estimate of 1 year as generation time and plotted on a linear timescale. We also used these demographies in calculating split times between subpopulations in a cross-coalescent framework of SMC++. The distinguished pairs were determined as described above, with the difference that each individual of the pair belonged to a different subpopulation.

**1.5. Archaeological and paleoenvironmental context**

Using a comprehensive database of rice archaeological records (*13*) in 1,000-year intervals, we plotted two-dimensional distributions (for latitude and longitude) using GEOM_DENSITY2D and STAT_DENSITY2D functions from GGPLOT (*94*) onto a map of Asia in R v.3.4.1 (*78*).

In order to predict how changing temperatures might have impacted distribution of different types of rice (indica, and temperate and tropical varieties of japonica) we used a global record of Holocene temperatures (*36*) to reconstruct growing degree-days (GDDs) following the methods of d’Alpoim and Bocinsky (*35*). We derived daily modern temperatures from Global Historical Climatology Network weather stations across East, South, and Central Asia (*99*). To account for spatial heterogeneity in how stations at different altitudes respond to climatic change, we used variance matching and modulated maximum and minimum mean weather station climatology by SDs derived from Marcott *et al*. (*36*). This was carried out for each year in the Marcott record. The niche of different types of landraces was established by thresholding annual GDDs, a measure of accumulated units of heat required by plants to complete their life cycle. We then used indicator kriging to spatially interpolate these niches across the ETOPO5 5 arc min (c. 10 km) resolution elevation model (*100*). The full research compendium that contains all the code and data necessary to reproduce this analysis is available at: <https://github.com/bocinsky/gutaker2019>.

**3. Supplementary figures**

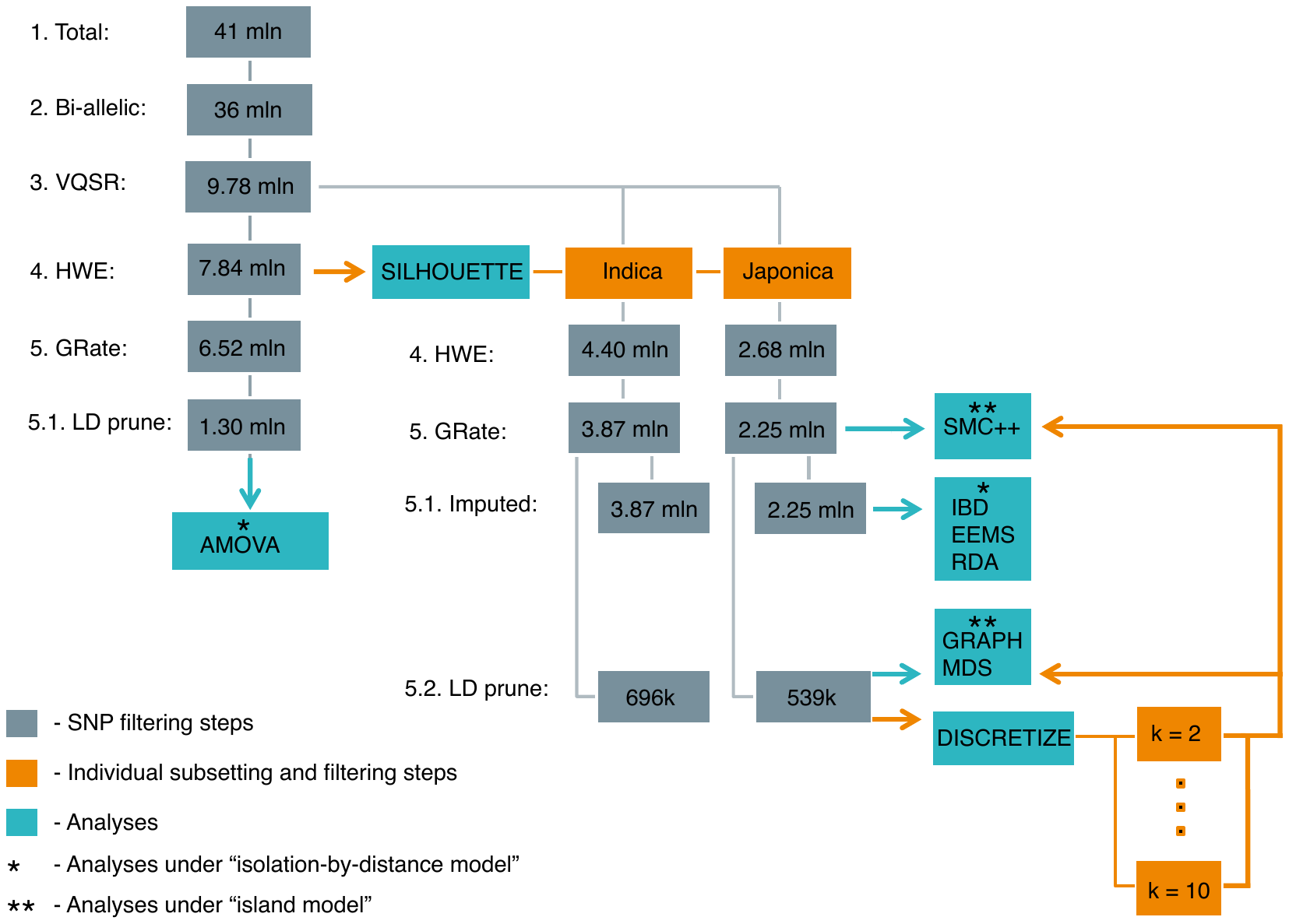

**Supplementary Fig. 1: The flowchart of data filters and analyses.** This flowchart tracks SNP and individual sets used for each analysis conducted in this study. Grey boxes represent numbers of SNPs retained after each filtering step (annotated with numbers 1-5). Orange boxes represent set of individuals used for filtering and analyses. Cyan boxes represent downstream analyses.

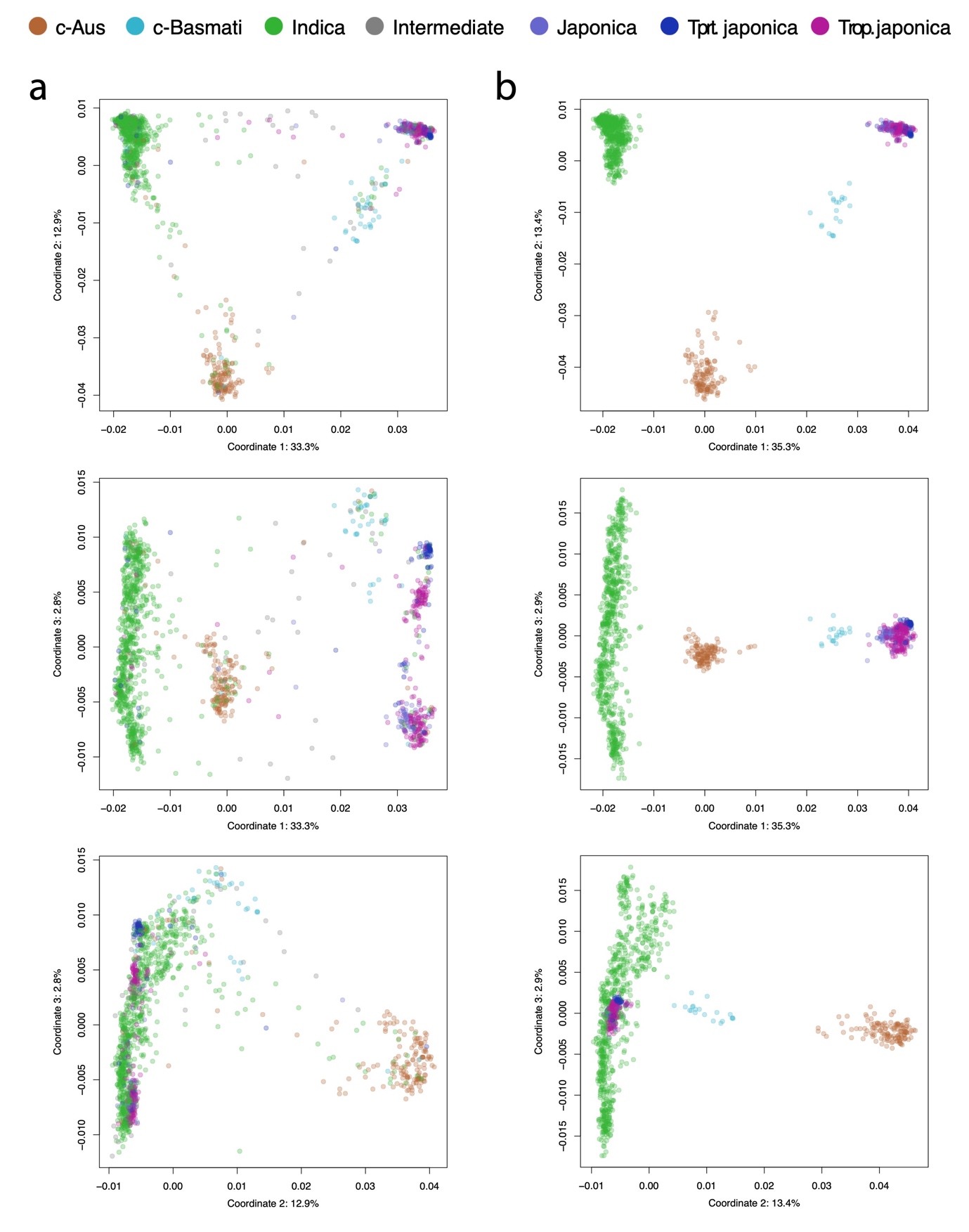

**Supplementary Fig. 2: Multidimensional scaling of all landraces.** Axes represent first, second and third dimension resultant from multidimensional scaling of genomic diversity for a total (**a**) and silhouette filtered (**b**) set of landraces. Each dimension is annotated with a proportion of genomic variation that it explains. Colors correspond to reported subgroup affiliation.

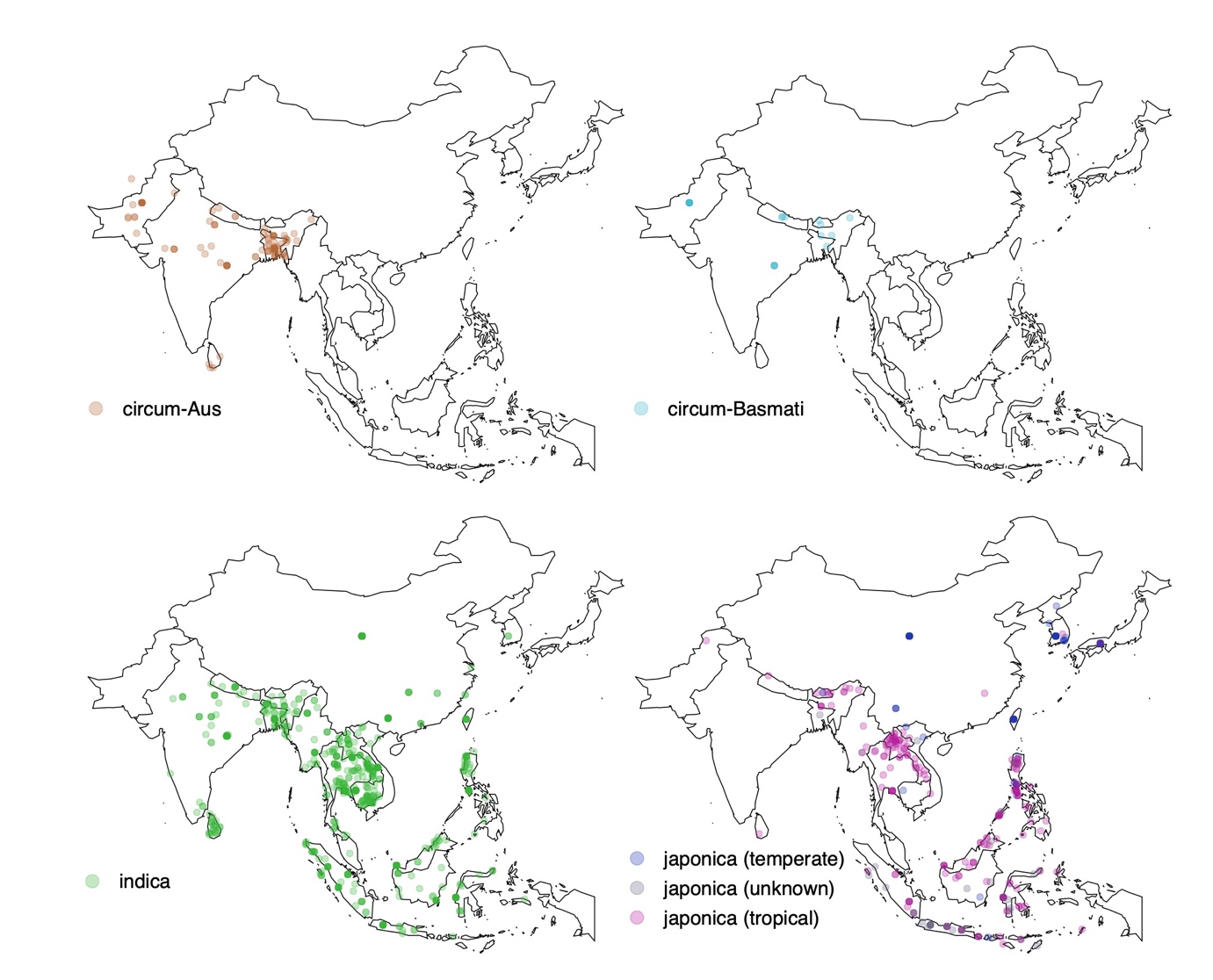

**Supplementary Fig. 3: Geographic origin of sampled genomes.** Maps of Asia with locations of sampled genomes for four subgroups: circum-Aus, circum-Basmati, indica, and japonica.

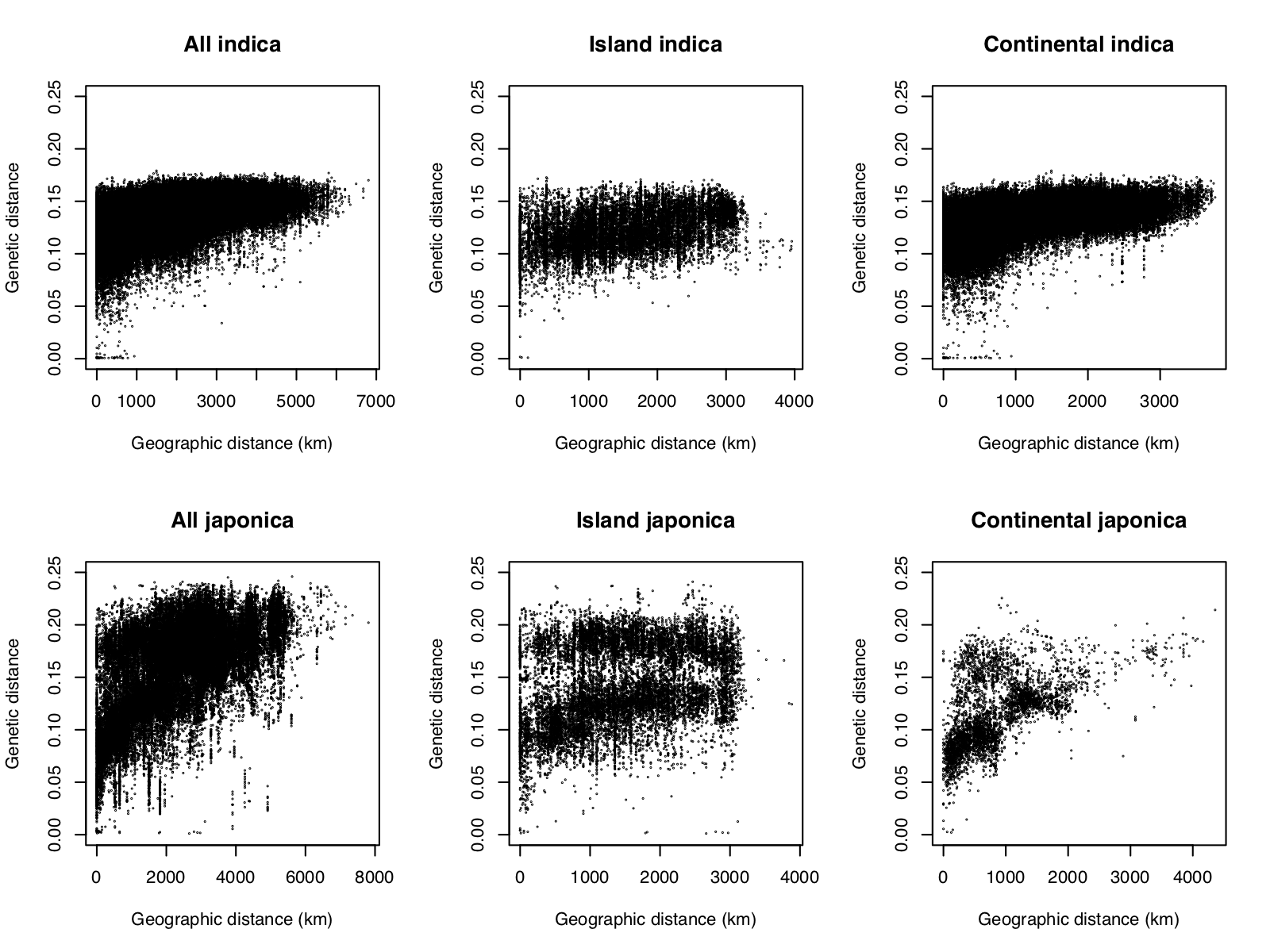

**Supplementary Fig. 4: Scatter plots of geographic vs. genetic distances.** Scatterplots demonstrating association between genetic and geographic distance in indica and japonica rice. For each subspecies, three different sets were plotted: total set, distances among island, and distances among continental population.

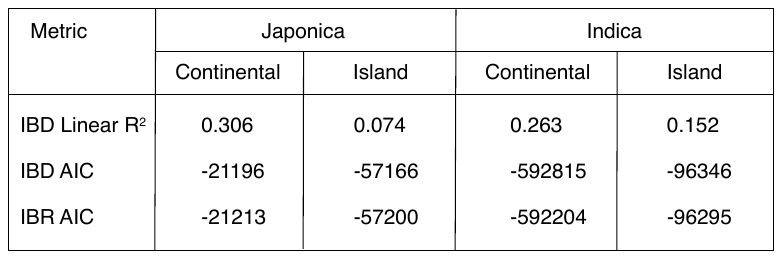

**Supplementary Fig. 5: Distance and travel time models.** Fit of isolation-by-distance (IBD) and isolation-by-resistance (IBR) models to japonica and indica in continental and island regions.

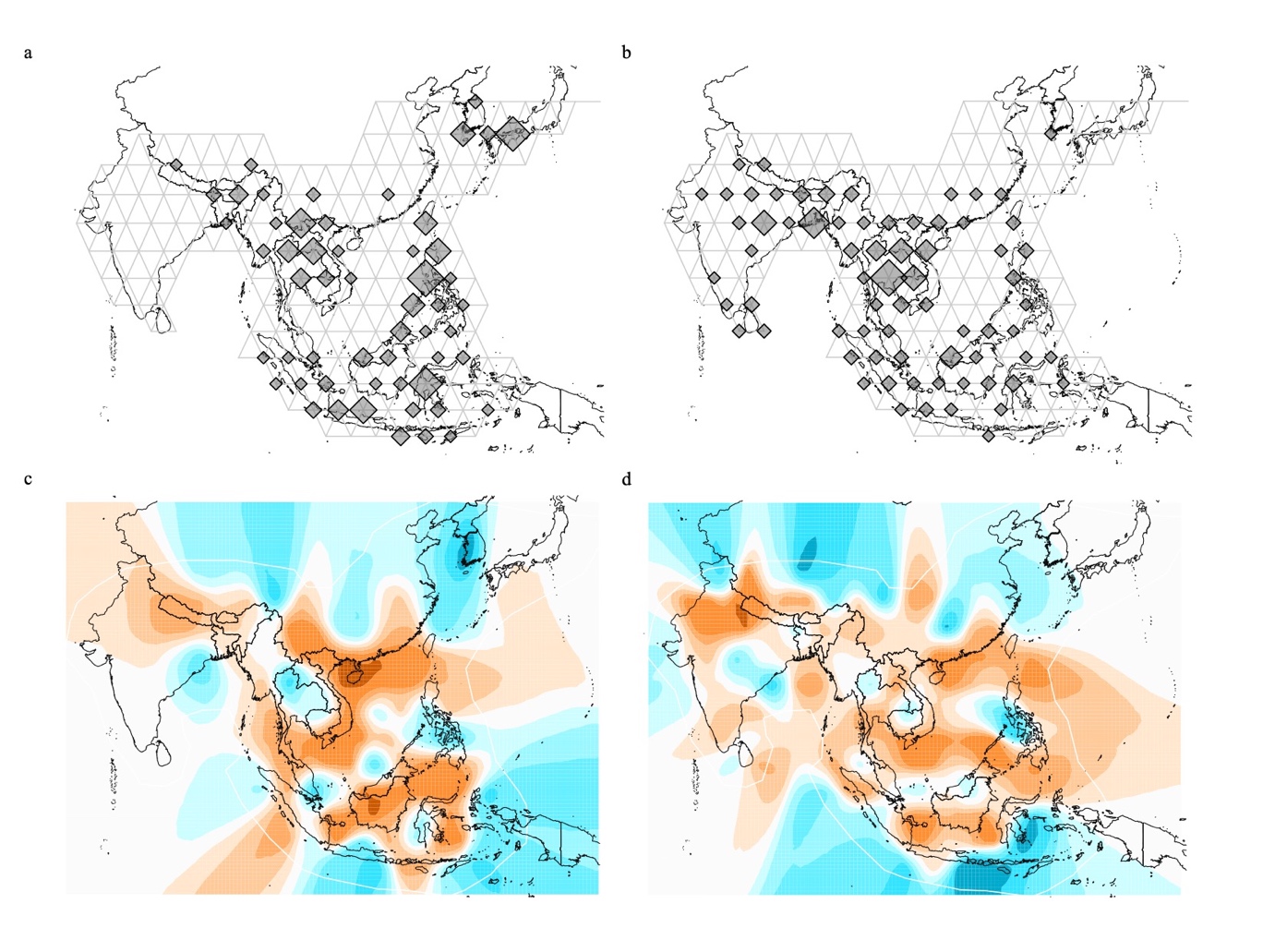

**Supplementary Fig. 6: Effective migration surfaces.** Map of demes on triangular grid for (**a**) japonica and (**b**) indica rice. Location of sample was used to assign each landrace into deme. The size of each deme is proportional to the number of landraces assigned. Effective migration surfaces from Vornoi tessellation of demes for (**c**) japonica and (**d**) indica rice. Orange colors represent areas of under-average migration with darker shades representing stronger barrier, while cyan colors represent areas of above-average migration.

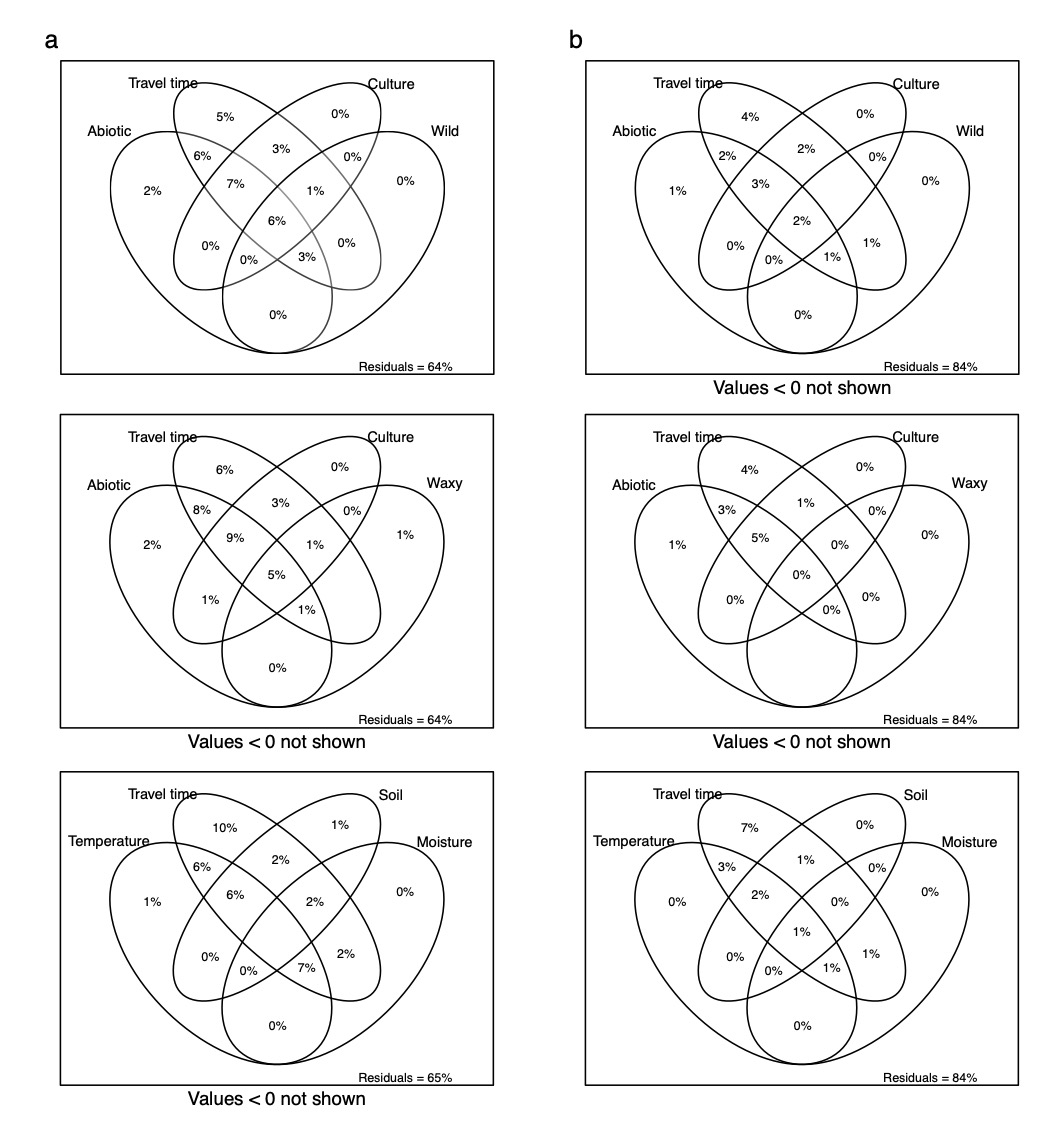

**Supplementary Fig. 7: Predictors of spatial distribution of genomic variation.** Four-way Ven diagrams for (**a**) japonica and (**b**) indica landraces representing the proportion (adjusted R^2^) of explained genomic variation.

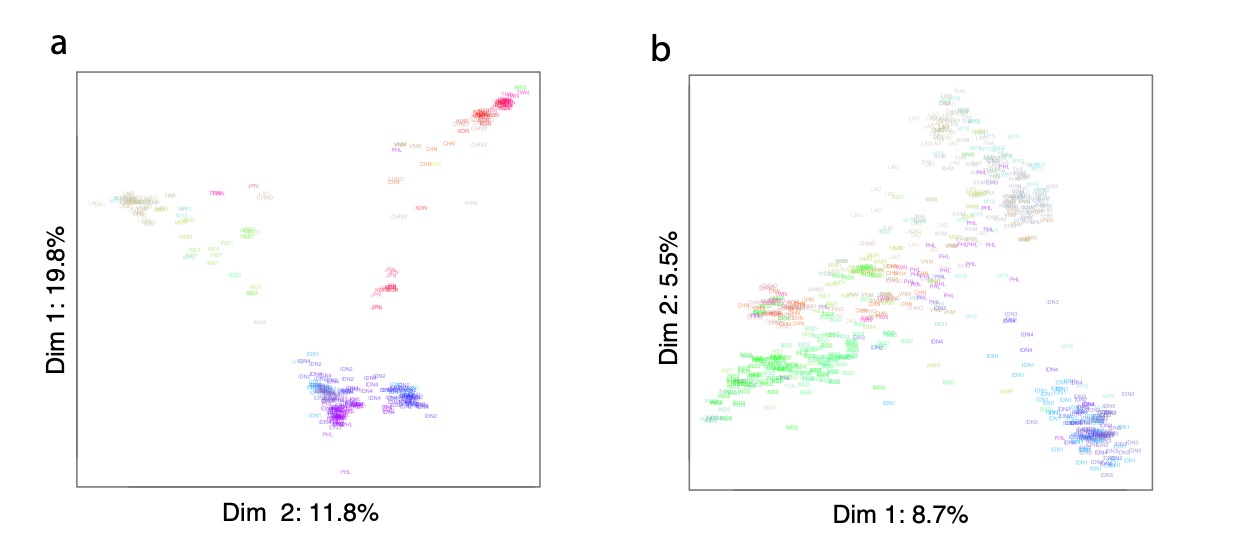

**Supplementary Fig. 8: Multidimensional scaling of japonica and indica rice.** Axes represent first and second dimension resulting from multidimensional scaling of genomic diversity for (**a**) japonica and (**b**) indica landraces. Each dimension is annotated with a proportion of genomic variation that it explains. Colors correspond to reported geographic regions of Asia.

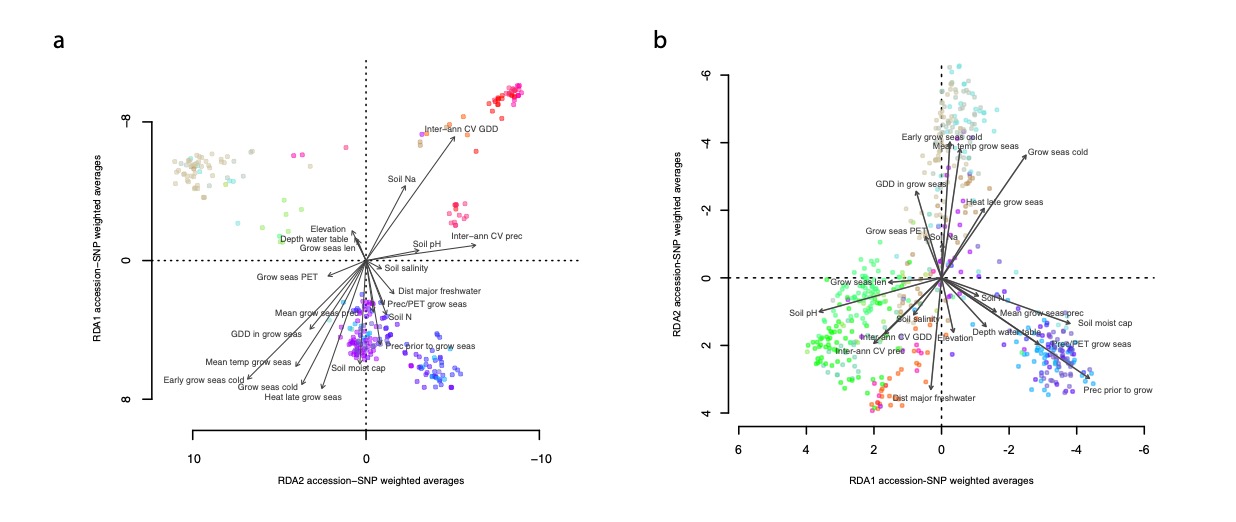

**Supplementary Fig. 9: Redundancy analysis with all variables.** (**a**) Japonica and (**b**) indica genotypes projected on the first two canonical axes of redundancy analysis. Arrows represent environmental predictors correlating with a maximal proportion of linear combinations of SNPs.

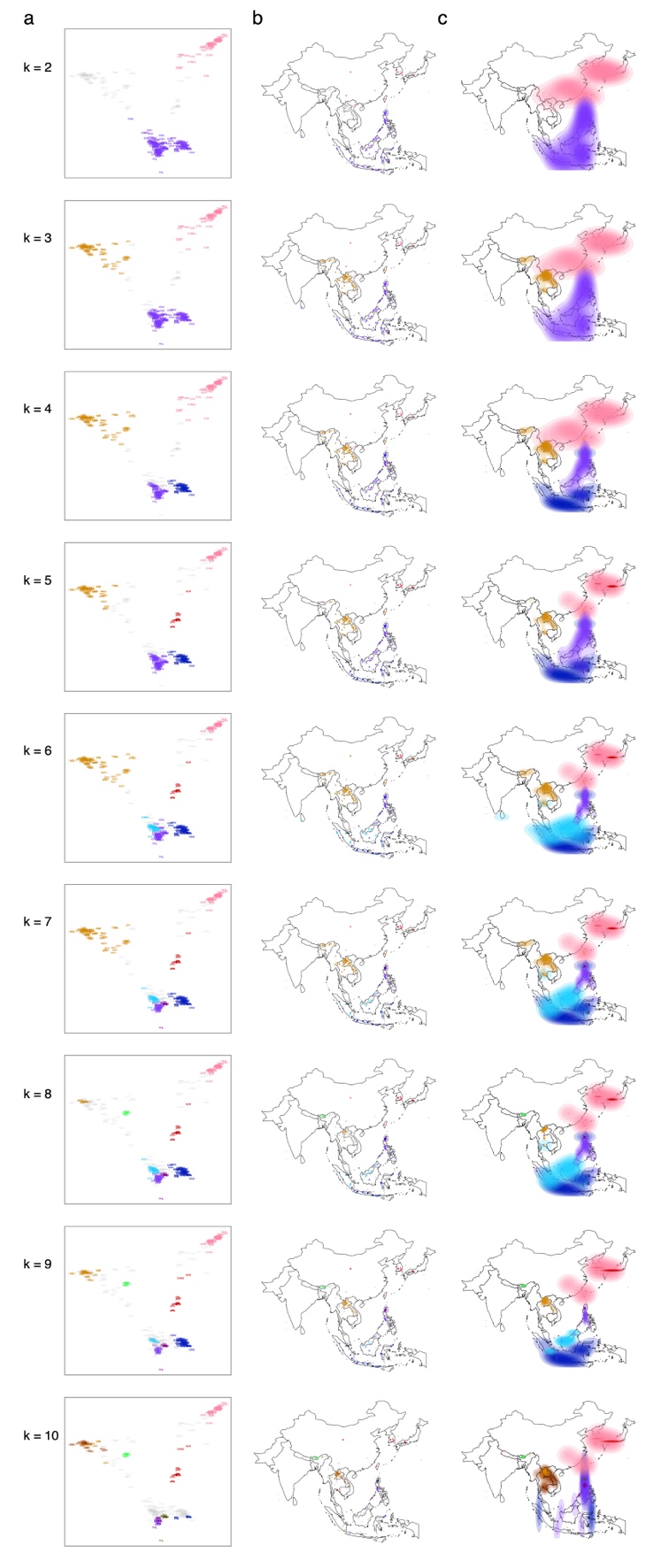

**Supplementary Fig. 10: K-medoids clustering of japonica genomes.** (**a**) Two first dimensions of multidimensional scaling of genomic diversity of japonica. K-medoid clusters after silhouette filtering are represented with different colors (number of populations, k ranges from 2 to 10). (**b**) Maps of distribution of landraces assigned to different discrete clusters are colored corresponding to k’s from 2 to 10. (**c**) The geographic distribution of subpopulations represented as colored two-dimensional Kernel density fields.

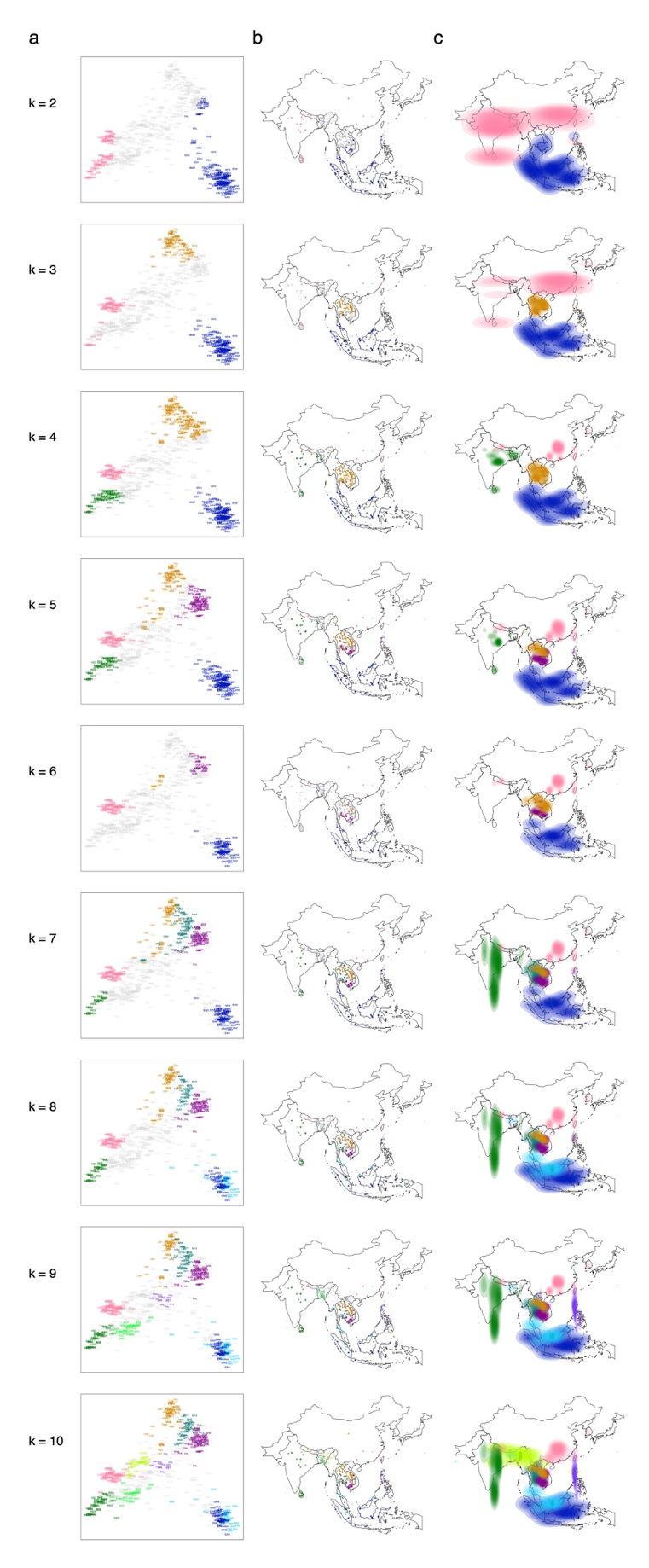

**Supplementary Fig. 11: K-medoids clustering of indica genomes.** (**a**) Two first dimensions of multidimensional scaling of genomic diversity of indica. K-medoid clusters after silhouette filtering are represented with different colors (number of clusters, k ranges from 2 to 10). (**b**) Maps of distribution of landraces assigned to different discrete clusters are colored corresponding to k’s from 2 to 10. (**c**) The geographic distribution of subpopulations represented as colored two-dimensional Kernel density fields.

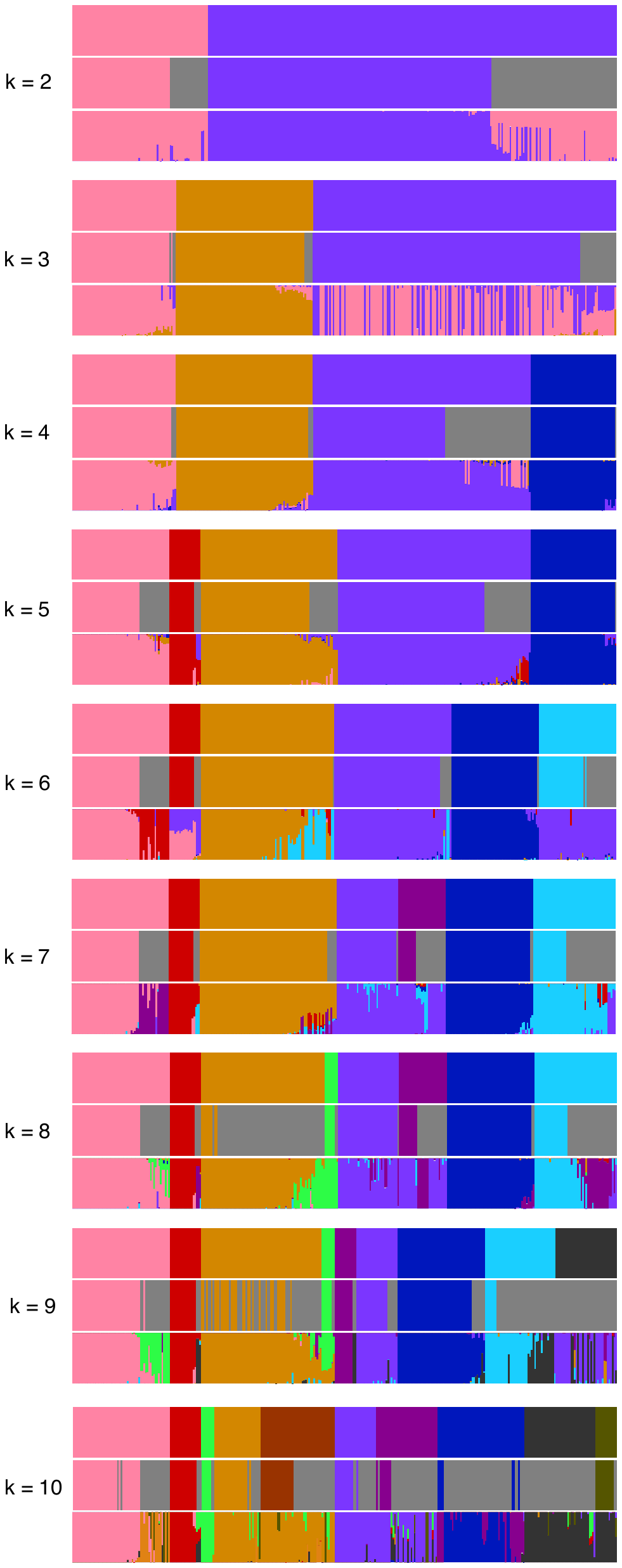

**Supplementary Fig. 12: Comparison of clustering approaches in japonica.** Analyses were carried out for number of japonica populations (k) from 2 to 10. For each k top panel represent the clustering by means of k-medoid algorithm, middle panel represent the result of ‘discretize’ filtering (grey), bottom panel shows ancestry proportions calculated with ADMIXTURE.

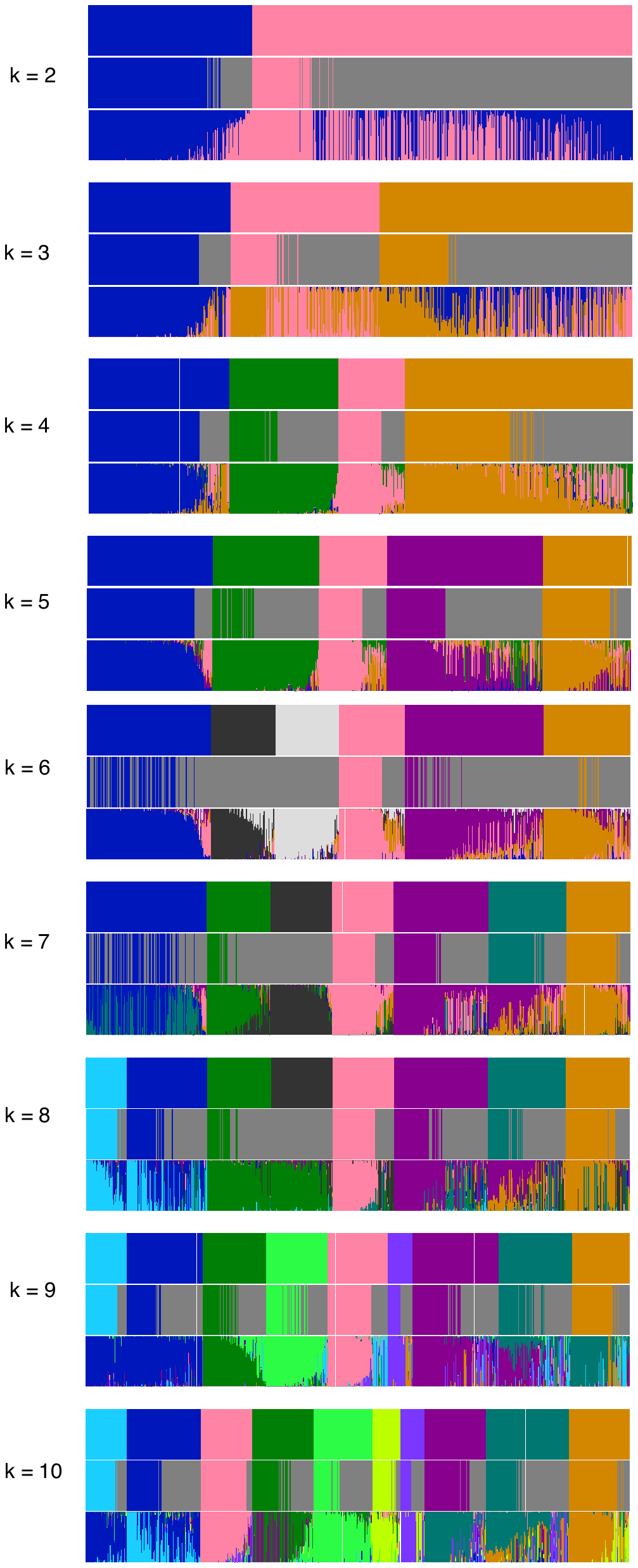

**Supplementary Fig. 13: Comparison of clustering approaches in indica.** Analyses were carried out for number of indica populations (k) from 2 to 10. For each k top panel represent the clustering by means of k-medoid algorithm, middle panel represent the result of ‘discretize’ filtering (grey), bottom panel shows ancestry proportions calculated with ADMIXTURE.

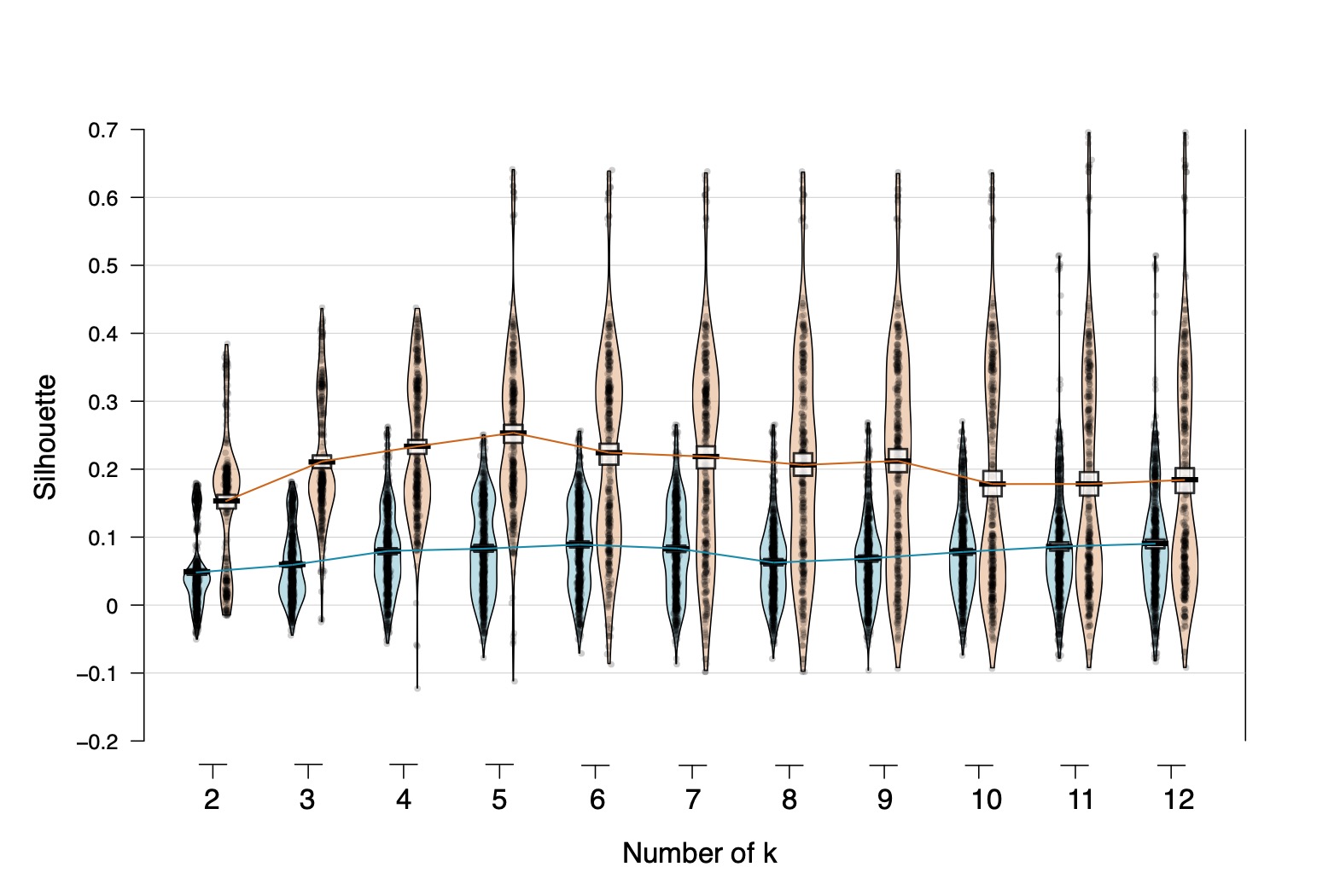

**Supplementary Fig. 14: Overview of silhouette scores.** Pirate plots of silhouette scores that were calculated for each landrace after k-medoid clustering in japonica and indica. Scores were calculated for k ranging from 2 to 12.

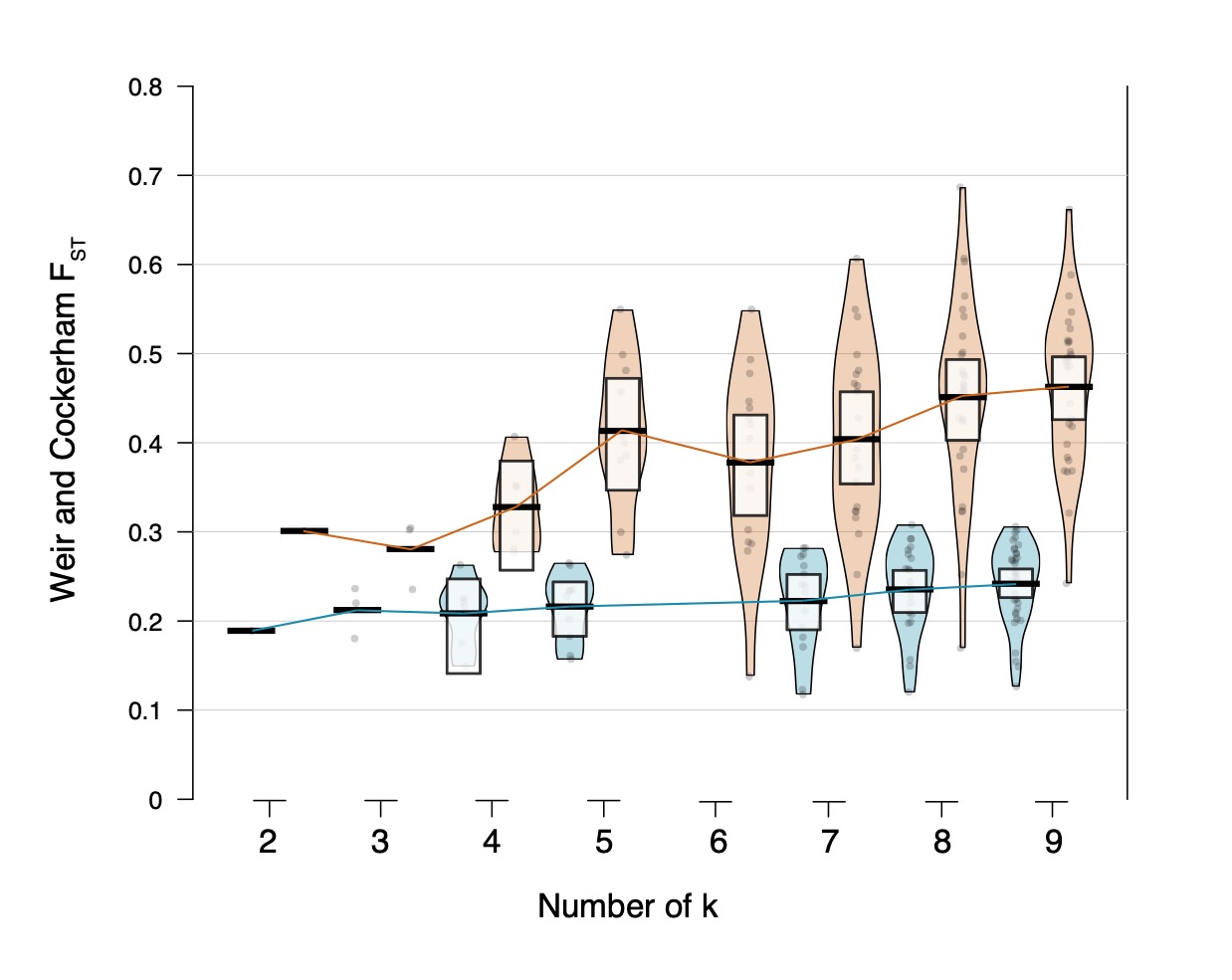

**Supplementary Fig. 15: Cluster differentiation with pairwise F_ST_.** Pirate plots of weighted average pairwise F_ST_ values after k-medoid clustering and ‘discretize’ filtering in japonica and indica. F_ST_ values were calculated for k ranging from 2 to 12.

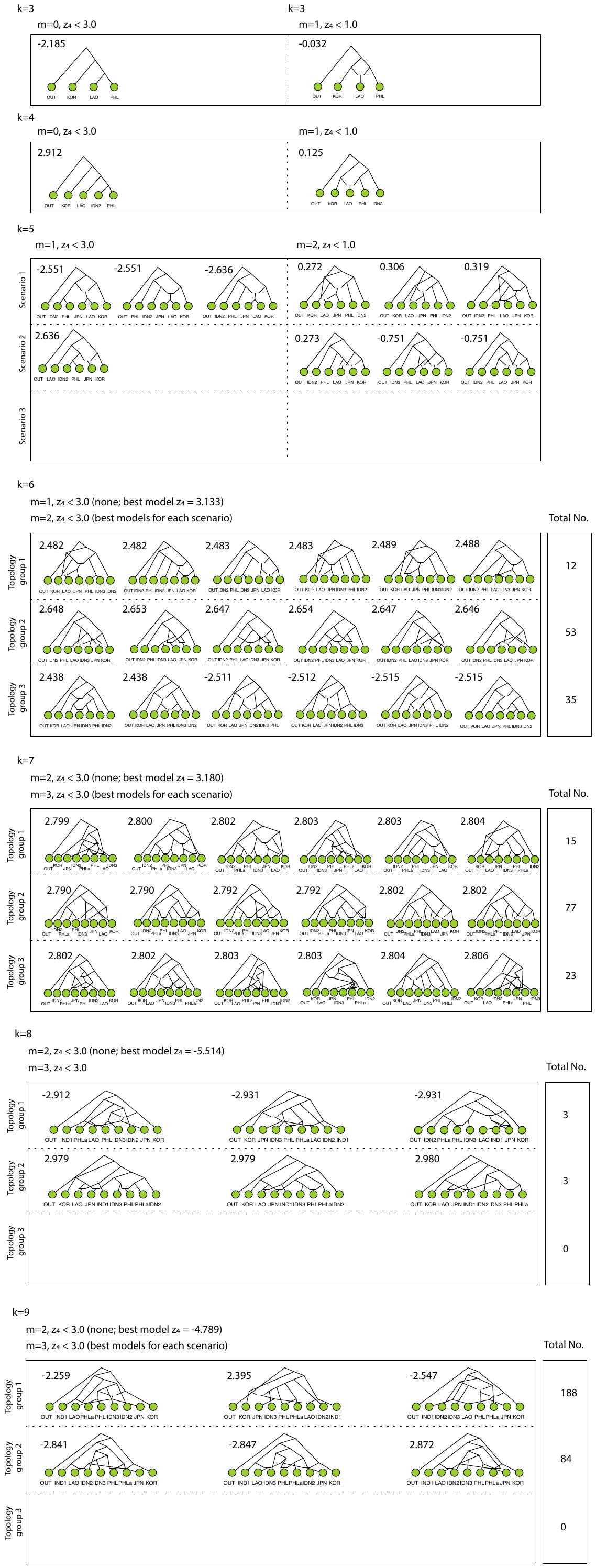

**Supplementary Fig. 16: Summary of Admixture Graphs in japonica.** Admixture graphs were summarized for number of populations, k ranging from 3 to 9 and number of admixture events, m ranging from 0 to 3. Values over admixture graphs represent worst-fit four-population drift z-score. Starting from k = 5, models were summarized into three different topology groups. Starting from k = 6, number of data-fitting alternative topologies were reported for each topology group.

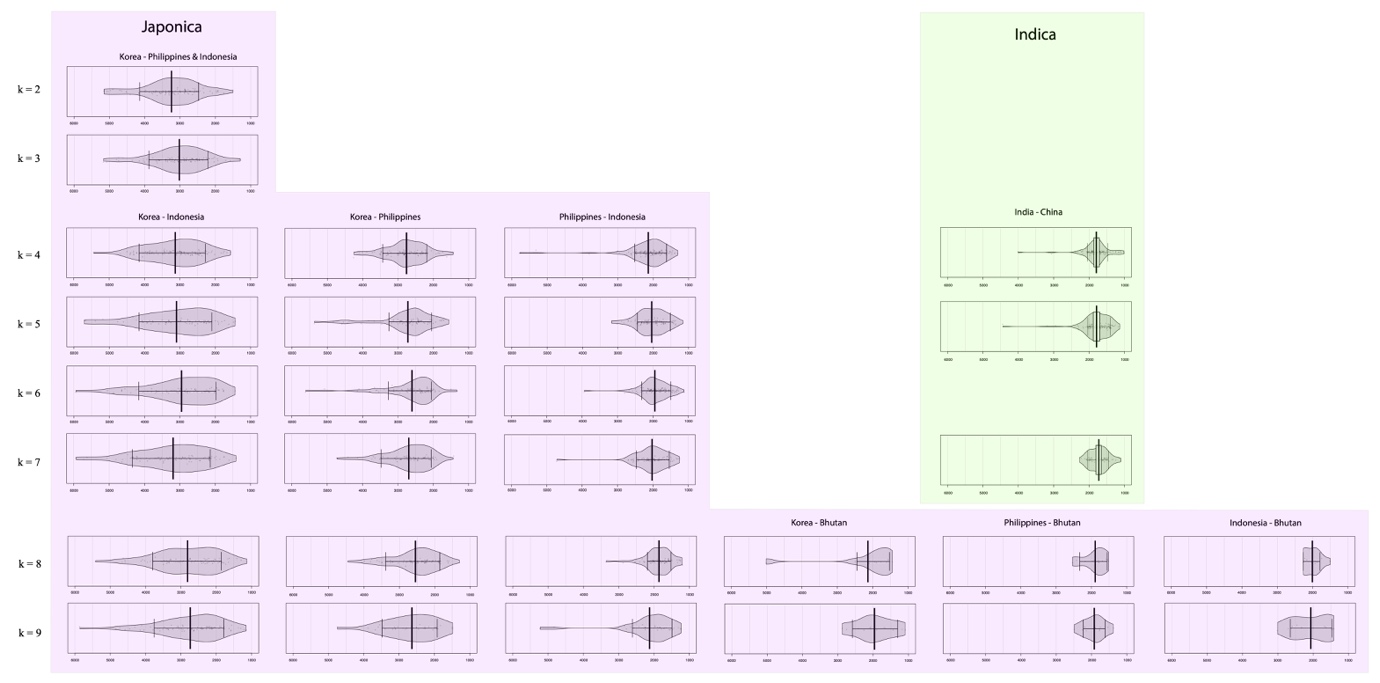

**Supplementary Fig. 17: Split times within japonica and indica.** Pirate plots representing split events for pairs of landraces from different two subpopulations in time (x-axis, linear scale) for number of subpopulations, k ranging from 2 to 9.

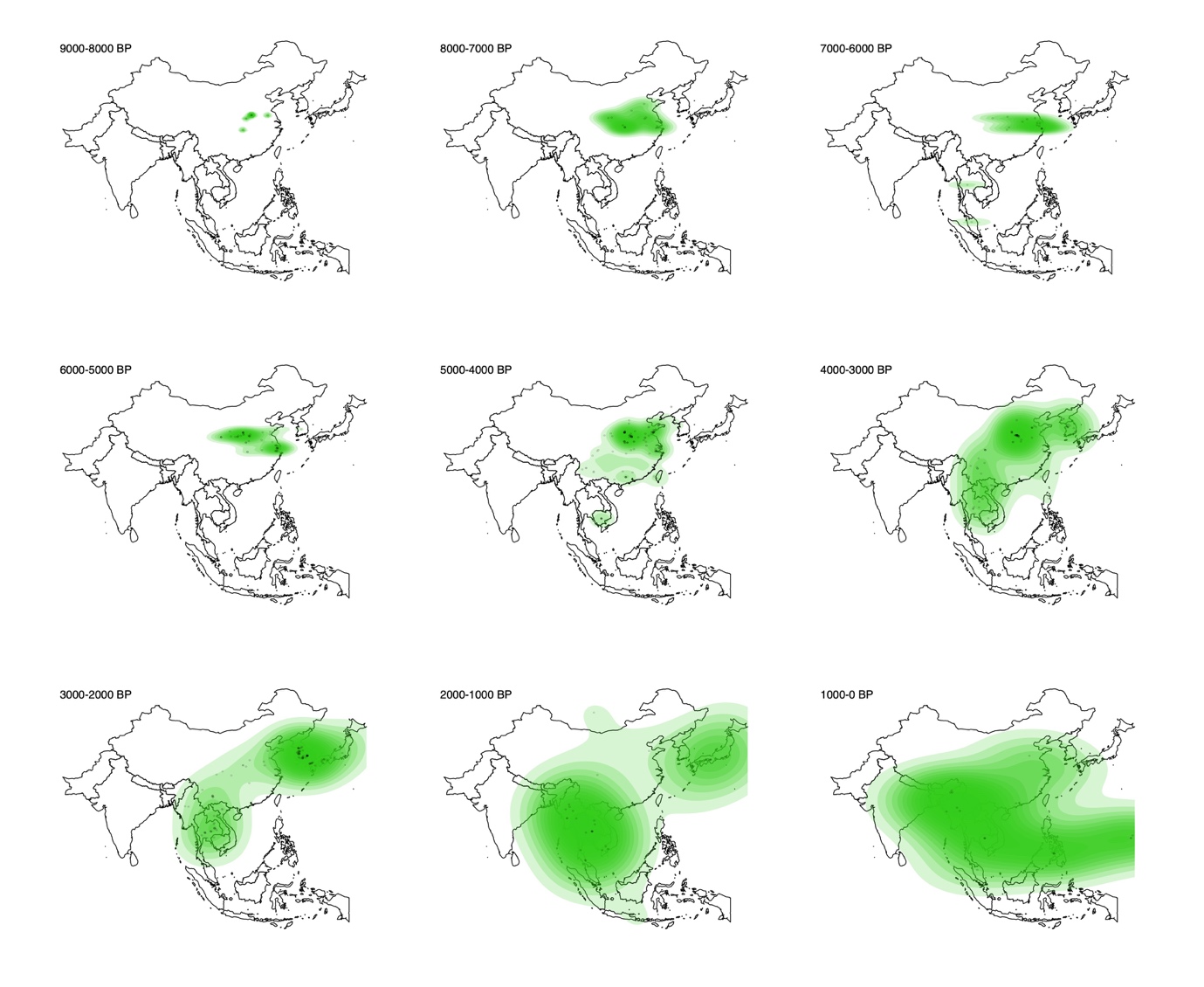

**Supplementary Fig. 18: Spatio-temporal distribution of rice archaeobotanical remains.** Maps of Asia in 1,000-year intervals annotated with archaeological records of rice cultivation. Their geographic distributions were represented as colored two-dimensional Kernel density fields.

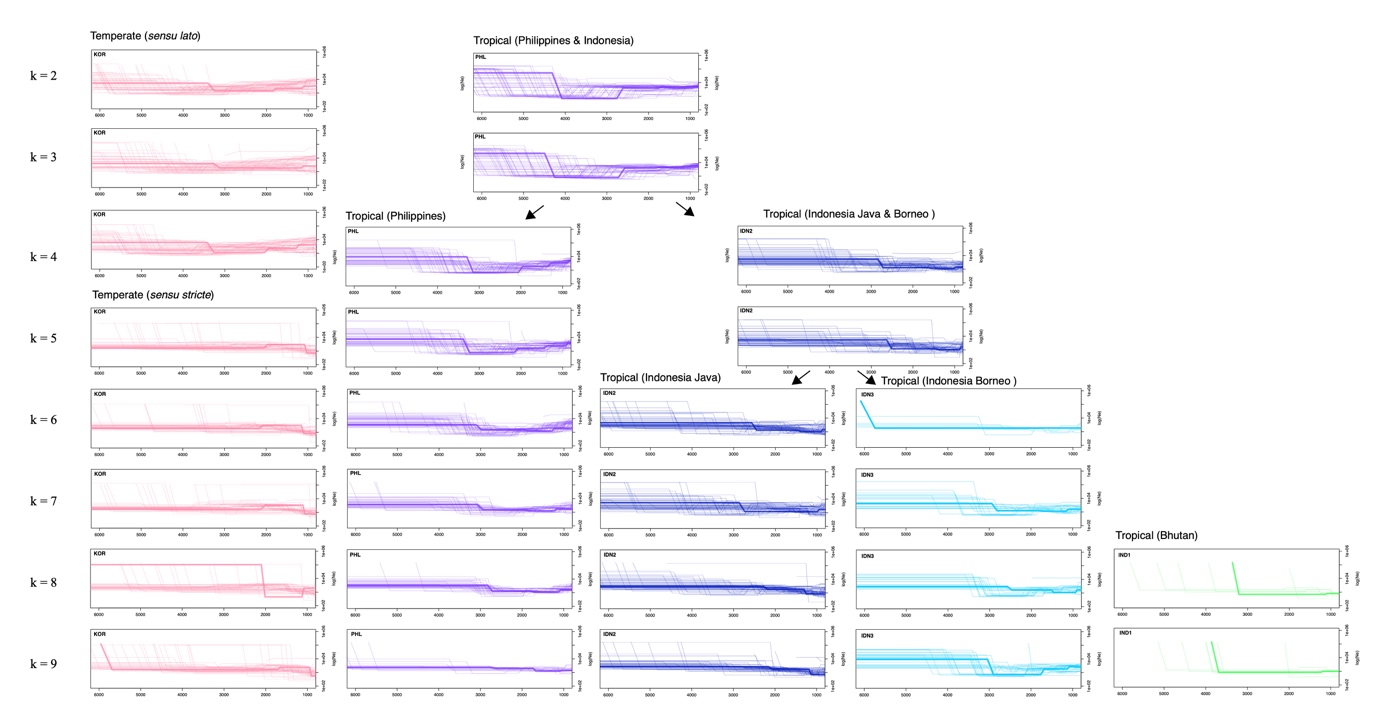

**Supplementary Fig. 19: Demographic histories in japonica subpopulations.** Plots represent changes in effective population sizes (y-axis, log scale) in time (x-axis, linear scale) of japonica subpopulations for number of populations, k between 2 and 9. Thin lines represent demographic model for a single landrace, while thick lines represent joint demographic models for all sampled landraces in each subpopulation.

**Supplementary Fig. 20: Spatio-temporal distribution of rice thermal niche.** Maps illustrating probability of rice being in niche based on minimum and maximum growing degree days requirement for temperate and tropical japonica landraces. Panels above represent probabilities before 4.2k event, 4410 years ago, while panels below represent probabilities after 4.2k cooling, 3510 years ago. White lines denote the border under which niche probability drops below 75%. Under each map is a plot of probabilities averaged across spatial scale. The mean (thick black line) and the interquartile range, 25% to 75% (gray shaded area) of probability of being in the thermal niche. The thin black lines are the mean probabilities using the lower and upper confidence intervals of the temperature reconstruction.

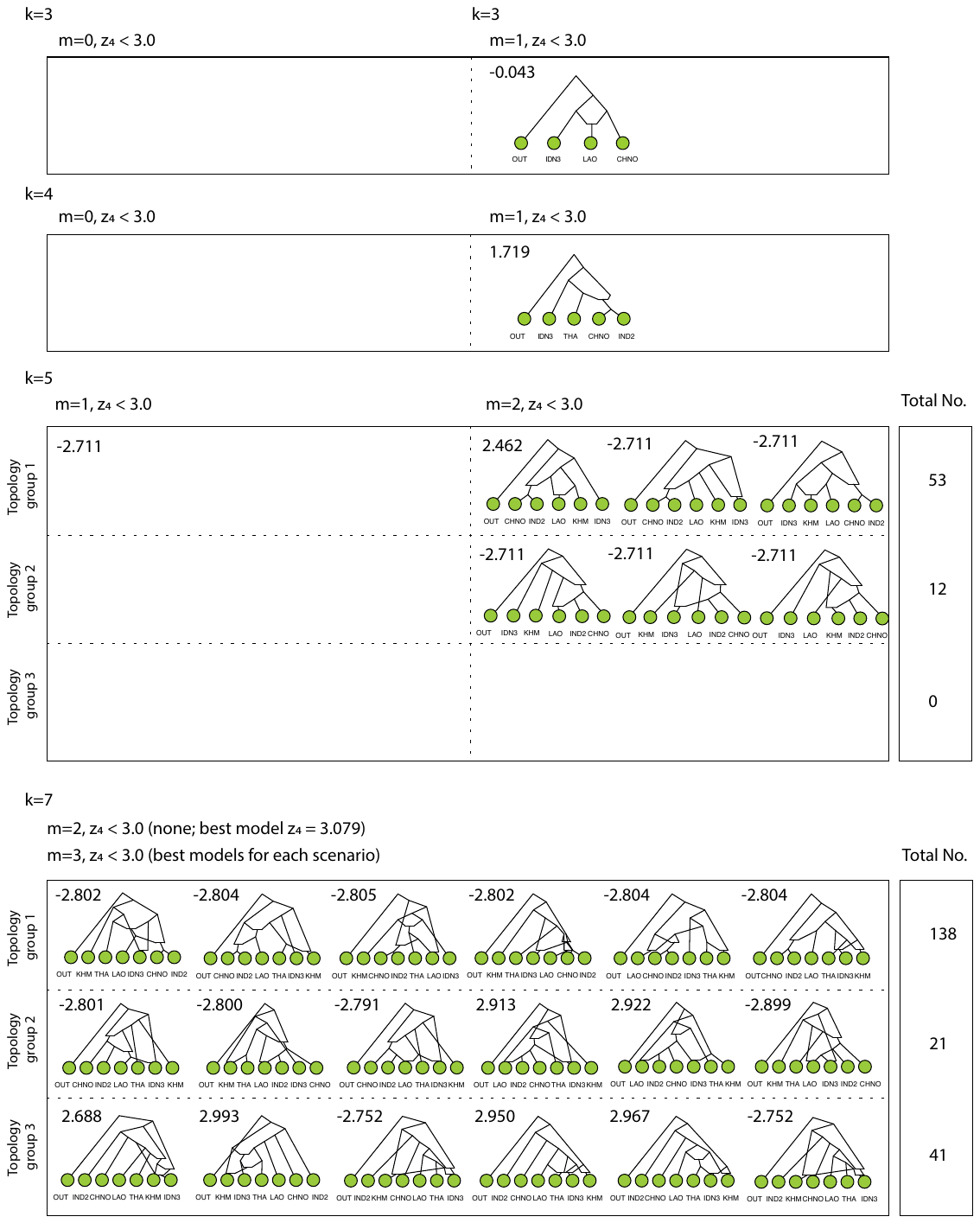

**Supplementary Fig. 21: Summary of Admixture Graphs in indica.** Admixture graphs were summarized for number of populations, k ranging from 3 to 7 and number of admixture events, m ranging from 0 to 3. Values over admixture graphs represent worst-fit four-population drift z-score. Starting from k = 5, models were summarized into three different topology groups, and number of data-fitting alternative topologies were reported for each topology group.

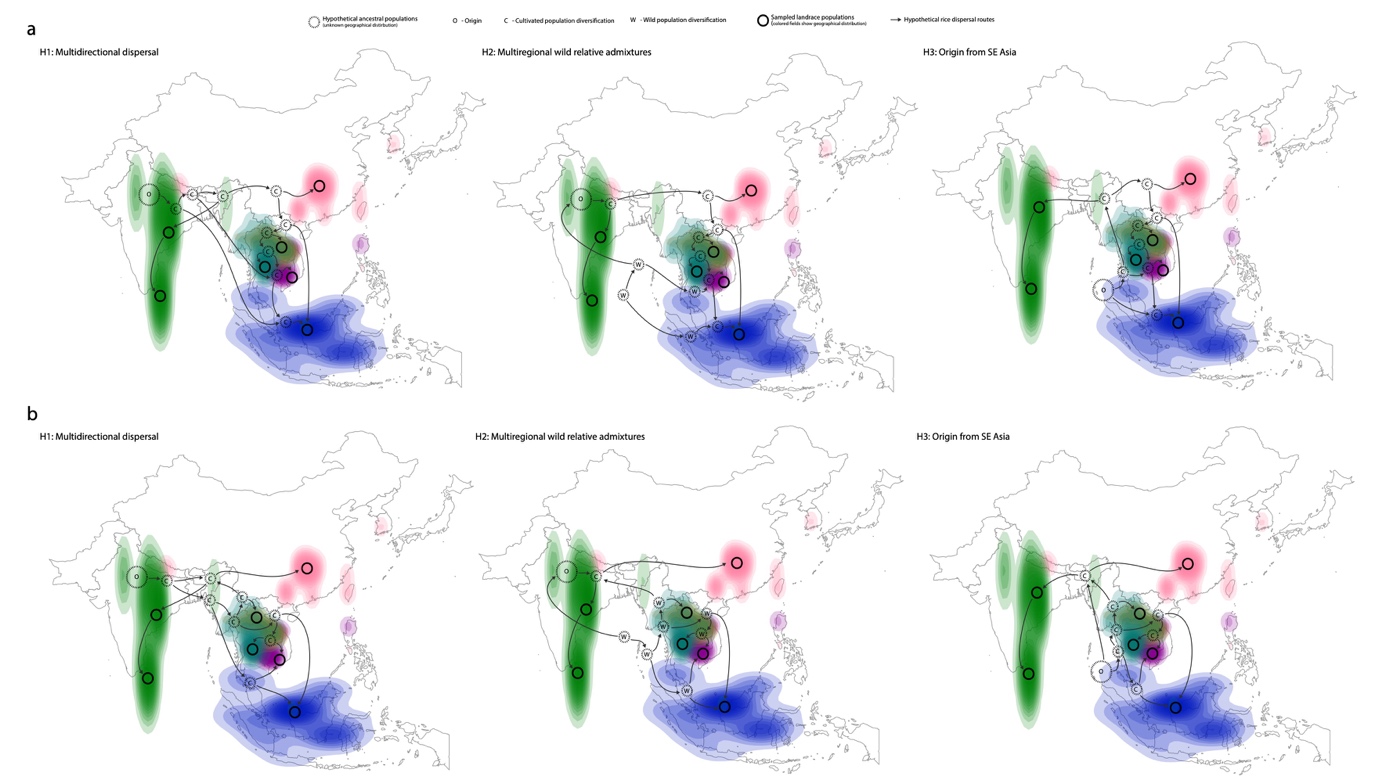

**Supplementary Fig. 22: Indica dispersal hypotheses.** Maps showing hypothetical indica dispersal in Asia based on (**a**) best admixture graph model at k = 7, and (**b**) admixture graph, of which topology was congruent with data between k = 2 and k = 7. Placement of internal nodes and directionality of dispersals captures admixture graph topologies in three scenarios: H1 - intense and multidirectional dispersal, H2 - dispersal followed by admixture with local wild relatives, and H3 - Southeast Asian origin of proto-indica.

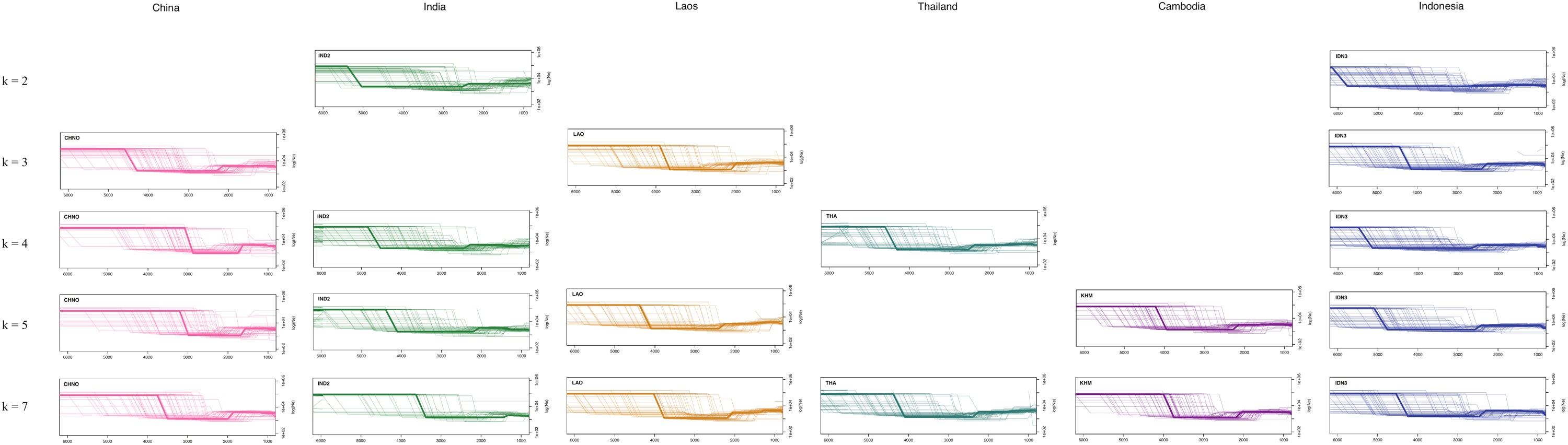

**Supplementary Fig. 23: Demographic histories in indica subpopulations.** Plots represent changes in effective population sizes (y-axis, log scale) in time (x-axis, linear scale) of indica subpopulations for number of populations, k between 2 and 7. Thin lines represent demographic model for a single landrace, while thick lines represent joint demographic models for all sampled landraces in each subpopulation.
